# Telomere Dysfunction in Human Astrocytes Drives Acrocentric Chromosome Instability and Nucleolar Reorganization

**DOI:** 10.64898/2026.05.19.726354

**Authors:** Ogechukwu Mbegbu, Yi-An Chen, Yue Hao, Noelle Fukushima, T. Rhyker Ranallo-Benavidez, Ava Bryan, Szehoi Chan, Nikki Messick, Maria Kyriakidou, Tianpeng Zhang, Pippa F. Cosper, Floris P. Barthel

## Abstract

Loss of p53 and Rb enables continued cell division despite progressive telomere erosion, ultimately triggering telomere crisis. While recent work has clarified mechanisms of chromatin bridge resolution, the longitudinal dynamics of structural damage through crisis remain incompletely understood. We transduced normal human astrocyte (NHA) cells with HPV18 E6/E7 and tracked them across extended population doublings. NHA E6/E7 cells showed progressive telomere shortening, anaphase bridges, and a growth plateau consistent with crisis. Multicolor FISH revealed subclonal chromosomal abnormalities largely invisible to short-read sequencing, with acrocentric chromosomes disproportionately affected: chromosome 13 was abnormal in over 92% of metaphases, and chromosomes 21 and 22 were similarly enriched. Translocations involving chromosome 13 were transient, replaced at later passages, while numerical aberrations persisted. Immunofluorescence revealed compact spherical nucleoli replaced by dispersed necklace-like structures, indicating a large-scale reorganization of nucleolar structure in response to telomere dysfunction. To determine whether these changes reflected altered chromosomal organization in the nucleus, we performed Hi-C and deployed KaryoScope, our alignment-free k-mer-based approach that recovers trans-chromosomal signal from repetitive acrocentric short arms. Inter-chromosomal contacts among nucleolar organizing region-bearing acrocentric chromosomes were markedly and persistently depleted in E6/E7 cells. Together, cytogenetic, imaging, and chromatin-contact data identify the nucleolus as a structural nexus linking telomere dysfunction to large-scale genomic rearrangement.

## Introduction

Genome stability in human cells depends critically on the p53 and Rb cell cycle checkpoints, which jointly enforce proliferative arrest in response to DNA damage [1]. When cells with critically short telomeres engage these checkpoints, senescence is triggered [2, 3]. Suppression of the p53 or Rb pathway allows cells to bypass these proliferative limits, driving continued division despite progressive telomere erosion and the accumulation of genomic aberrations [4, 5]. The resulting period of telomere dysfunction, characterized by breakage-fusion-bridge (BFB) cycles, end-to-end chromosome fusions, and catastrophic rearrangements, culminates in telomere crisis, the acute growth arrest that represents a critical bottleneck in tumor evolution [6–8]. Recent mechanistic studies have defined specific mechanisms of chromatin bridge resolution, including nucleolytic processing by TREX1 and mechanical disruption of the nuclear envelope [9, 10]. However, longitudinal tracking of structural variants through telomere crisis revealed a complexity of genomic rearrangements that exceeded what these individual mechanisms predicted [11]. Which chromosomes are affected, what rearrangements arise, and how nuclear organization is disrupted remain open questions.

Not all chromosomes are equally vulnerable to rearrangements [12]. The five human acrocentric chromosomes (13, 14, 15, 21 and 22) share a unique genomic architecture. Their short arms harbor nucleolar organizing regions (NORs) containing ribosomal DNA (rDNA) arrays that coalesce each cell cycle to form the nucleolus [13, 14]. The rDNA arrays themselves are among the most unstable loci in eukaryotic genomes, with copy-number variation and recombination documented across yeast, Drosophila, and human contexts including aging, cancer, and neurodegeneration [15]. Acrocentric chromosomes thus combine spatial clustering at the nucleolus with intrinsic instability of their NOR-bearing short arms, two features that may converge to make them preferred partners for inter-chromosomal fusions when telomeres become deprotected. Whether this vulnerability extends to progressive telomere erosion in p53/Rb-impaired cells in a prolonged state of telomere dysfunction has not been tested.

To capture the longitudinal arc of telomere dysfunction from senescence bypass through telomere crisis, we adapted the established NHA E6/E7 system without hTERT and tracked primary human astrocytes across extended population doublings [16, 17]. This window is bypassed entirely in hTERT-stabilized models [8] and only briefly accessed in acute telomere deprotection experiments [18, 19], leaving the longitudinal accumulation of structural variants in primary diploid cells largely unexplored. We found that telomere dysfunction in p53/Rb-impaired astrocytes gives rise to numerical and structural chromosomal aberrations, and use longitudinal sampling to reveal how these aberrations accumulate and evolve over time. Remarkably, preferential vulnerability of acrocentric chromosomes was accompanied by disruption of nucleolar architecture, pointing to a previously unappreciated link between telomere dysfunction and nucleolar biology.

## Results

### NHA E6/E7 cells recapitulate a telomere crisis phenotype

The NHA-E6/E7 system was originally developed as the basis for a glioma transformation model, in which HPV E6/E7 was co-expressed with hTERT and oncogenic Ras to derive an isogenic cell line that would grow tumors in laboratory animals [16, 17]. However, the simultaneous expression of hTERT circumvented telomere shortening and the genomic consequences of that process that may be characteristic of these tumors [20–22]. To faithfully capture the period of telomere dysfunction that is bypassed by early telomerase activation, we independently transduced primary NHA cells with HPV18 E6/E7 or hTERT (Figure 1A).

**Figure 1:**
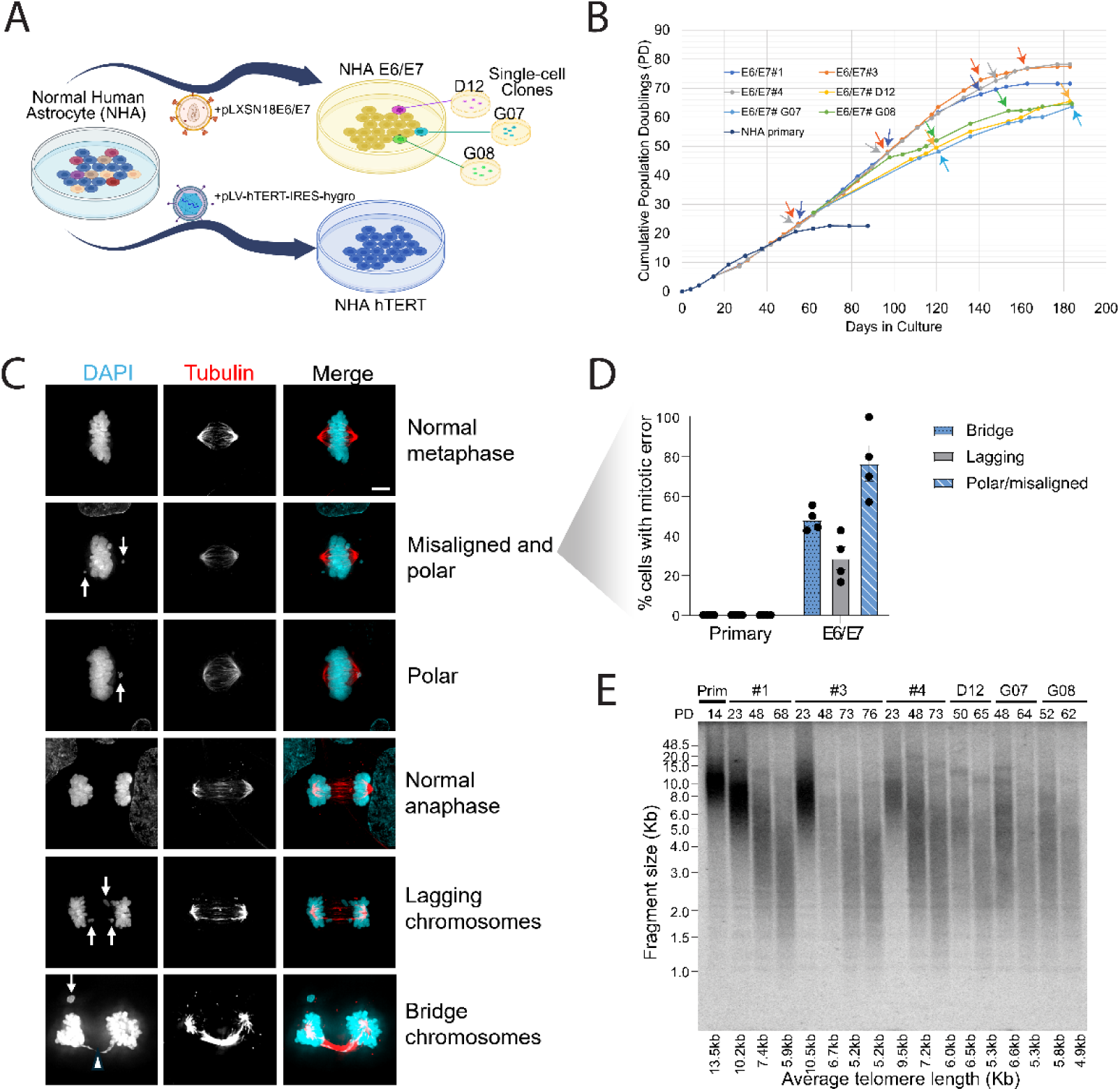
HPV18 E6/E7 expression promotes immortalization and genomic alterations in normal human astrocytes. **(A)** Schematic of the experimental workflow. Primary normal human astrocyte (NHA) cells were transduced with lentivirus to express either hTERT or HPV18 E6/E7. Single-cell clones were selected from E6/E7 bulk transduced cells. (**B)** Growth curves of primary NHA cells, E6/E7-expressing bulk cultures (E6/E7 bulk #1, #3, #4), and E6/E7-expressing single-cell clones (D12, G07, G08). Cumulative population doublings (PDs) are plotted over time. Circles indicate collection points, and arrows denote time points used for DNA sequencing. **(C)** Immunofluorescence examples of NHA E6/E7 metaphase or anaphase cells with mitotic errors. The image shows arrows highlighting misaligned/polar and lagging chromosomes and an arrowhead indicating bridge chromosomes. Scale bar= 10 □m. **(D)** Quantification of CIN in n=4 biological replicates of NHA primary and E6/E7 expressing NHA cells. The mitotic index was very low for primary (0.4%) and E6/E7 cells (1.4%). For primary, a total of 16 cells in metaphase and 32 cells in anaphase/telophase are represented. For the E6/E7 expressing cells, an average of 12 cells in metaphase and 8 cells in anaphase/telophase were quantified for each biological replicate. **(E)** TRF analysis of telomere length in NHA primary and six E6/E7 clones at increasing PDs. Average fragment lengths for each sample are shown on the x-axis.

We monitored the growth characteristics of our engineered astrocytes over many successive population doublings (PDs). Primary NHA cells showed stagnated growth curves and positive staining for the senescence marker β-galactosidase after approximately PD 20-30, consistent with senescence (Figure 1B & Figure S1A-B). In contrast, NHA-hTERT cells continued growing well beyond this point and remained negative for β-galactosidase, consistent with immortal growth (Figure S1A-B). On the other hand, E6/E7-expressing bulk cultures (E6/E7 bulk #1, #3, #4) demonstrated significantly extended lifespans, continuing to proliferate beyond 180 days and reaching PD 70-80. β-galactosidase staining was not observed at high PD in NHA E6/E7 cells, indicating that a senescence response was successfully abrogated, likely via E6/E7-induced suppression of p53 as detected via Western blotting (Figure S1C-D). Single-cell clones (NHA E6/E7 clone D12, G07 and G08) derived from bulk culture #3 similarly displayed robust and sustained proliferation, with all clones surpassing PD 60. However, unlike NHA-hTERT transduced cells, NHA E6/E7 cells did not reach an immortal growth phase and all E6/E7 bulk and single cell clones demonstrated a second growth plateau at PD 60-80, consistent with severe ongoing telomere crisis and an equilibrium between cell division and death rather than senescence-induced growth arrest.

To verify that NHA E6/E7 cells were subject to telomere dysfunction prior to the crisis growth plateau, we utilized immunofluorescence staining to visualize and quantify mitotic errors (Figure 1C-D). Briefly, anti-tubulin antibodies were used to visualize spindles in mitotic cells during metaphase and anaphase, with chromosomes visualized by DAPI. Anaphase bridges, a hallmark feature of telomere dysfunction, were frequently observed in our E6/E7 cultures, and were not present in the primary cells. Moreover, other mitotic errors such as polar and lagging chromosomes were common in NHA E6/E7 cells compared to primary cells, reflecting telomere-independent chromosomal instability characteristic of HPV E6/E7 (Figure 1C-D) [23].

To confirm that NHA E6/E7 telomeres were shortening with increasing PD, we performed Southern blotting of telomeric terminal restriction fragments (TRF) using DNA extracts from primary NHA and E6/E7 cells at different PD (Figure 1E). As expected, TRF analysis showed a steady decrease in mean telomere length as PD increased, consistent with telomere shortening. Several NHA E6/E7 samples show a high molecular weight band, which was previously shown to be a marker for telomere fusions [24]. Taken together, these results are consistent with our NHA E6/E7 model showing varying degrees of telomere dysfunction, culminating in telomere crisis, a state in which progressive telomere shortening in the absence of an (immortality-granting) maintenance mechanism, produces critically short and deprotected chromosome ends that fuse and seed breakage-fusion-bridge (BFB) cycles, driving anaphase bridges, chromosome missegregation and pervasive mitotic catastrophe.

### Genomic instability and clonal evolution of NHA E6/E7 clones over prolonged culture

In order to evaluate the structural dynamics of NHA E6/E7 cells over time, we performed short read Illumina sequencing followed by analysis of copy number variation (CNVs), single nucleotide variants (SNVs) and small indels across different PDs (Figure 2).

**Figure 2:**
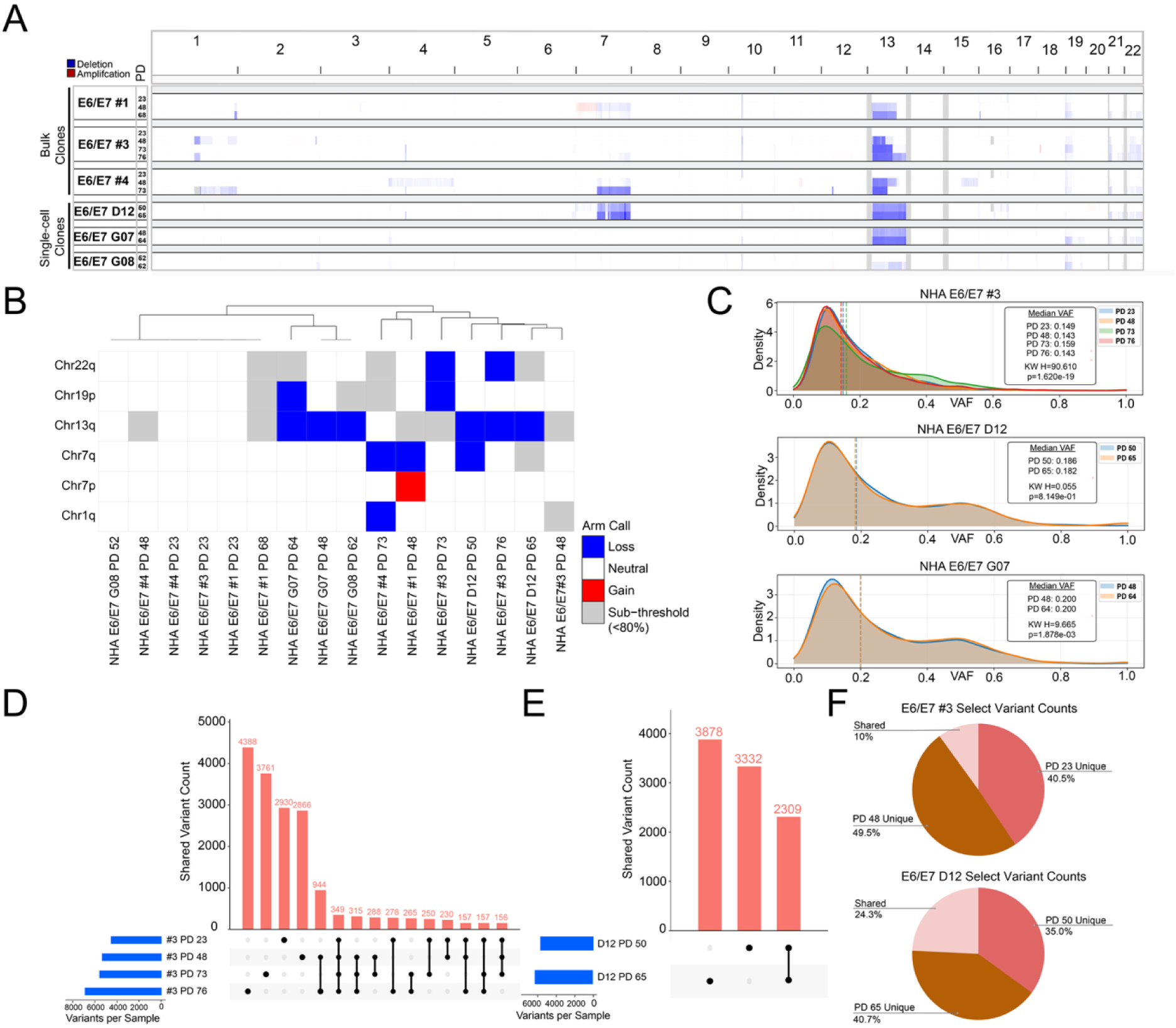
Genomic landscape and mutational evolution of E6/E7 clones. **(A)** Genome-wide CNV profiles for NHA E6/E7 bulk and single-cell clones across varying PDs. Deletions are indicated in blue and amplifications in red. Chromosome 13 deletion was observed in all but one of the clones sequenced. Chromosome 7 was also deleted in three of the clones. **(B)** Heatmap and hierarchical clustering of arm-level aneuploidy across samples. The color scale denotes chromosomal loss (blue), gain (red), neutral state (white), and sub-threshold alterations (<80%, grey). **(C)** Variant allele frequency (VAF) density distributions for representative samples (bulk #3, D12 and G07 clones) at progressive PDs. Dotted vertical lines denote median VAFs, with Kruskal-Wallis (KW) test statistics provided for each clone. **(D, E)** UpSet plots illustrating the intersection of shared and unique SNVs and indels across multiple PDs for bulk clone #3 **(D)** and single-cell clone D12 **(E)**.**(F)** Pie chart of selected variant counts for clone #3 and D12. PD 23 and 48 unique counts for clone #3 were plotted with their shared sum. PD 50 and 65 unique counts for clone D12 were also plotted with their shared sum.

Genome-wide CNV profiling of both bulk and single-cell clones revealed an accumulation of copy number changes over extended population doublings (Figure 2A, Supplementary Figure 2A). Most notably, a large-scale deletion on chromosome 13 was consistently observed across nearly all sequenced clones regardless of PD. Specifically, we observed that this anomaly often presented as a partial deletion at earlier PDs and progressed to a full deletion of chromosome 13 at later PDs. Furthermore, partial or complete deletions on chromosome 7 were observed in a specific subset of the clones. Hierarchical clustering of arm-level aneuploidies further localized these structural anomalies, confirming widespread loss of the 13q arm across most lineages, alongside more localized, clone-specific events such as 7q and 19p losses (Figure 2B).

To understand subclonal dynamics within these cell populations, we tracked variant allele frequency (VAF) distributions over time (Figure 2C). Density distributions for representative bulk (NHA E6/E7 #3) and single-cell (NHA E6/E7 D12 and G07) clones revealed complex shifting in mutational architectures. Specifically, within bulk clone #3, the largest median VAF was observed at PD 73; however, by PD 76, the median VAF reverted to levels comparable to earlier passages. Kruskal-Wallis statistical testing confirmed that the median VAFs and overall distributions shifted significantly (p < 0.05) across progressive PDs, most notably in bulk clone #3. These transient fluctuations suggest that rather than strict clonal selection, the subclonal architecture of NHA E6/E7 cells is heavily influenced by genetic drift and ongoing population remodeling, likely through the ongoing mitotic errors shown in Figure 1. Even though single-cell clones D12 and G07 did not exhibit obvious VAF shifts over time, compared to the bulk population, they displayed a distinct, elevated second peak in VAF around 0.5. This elevated peak was exclusively observed in the single-cell clones and absent in the bulk population, likely reflecting the expected fixation of heterozygous mutations characteristic of clonally derived lines.

We further quantified this continuous genomic evolution by mapping the intersection of shared and unique SNVs/indels across successive PDs (Figure 2D-F, Supplementary Figure 2B-E). UpSet plots demonstrated that while a small core of variants is maintained across all evaluated time points, cells continuously acquired new variants that did not persist (Figure 2D-E). For instance, in bulk clone #3, less than 400 variants were conserved from PD 23 through PD 76, yet thousands of unique variants distinctly emerged or disappeared at intermediate passages. Pie charts comparing selected early and late passages for bulk clone #3 and single-cell clone D12 similarly summarized this dynamic (Figure 2F). While a small number of mutations were shared between time points (e.g., 10% shared between PD 23 and 48 in clone #3), the majority of variants were passage-specific, underscoring continuous divergence and genomic instability of our NHA E6/E7 model over prolonged culture.

We further evaluated genomic stability over time by analyzing additional VAF distributions and mutational turnover (Supplementary Figure 2). While the 96-channel single-base substitution (SBS) spectrum for bulk clone #3 maintained a relatively consistent overall profile, absolute mutation counts exhibited fluctuations between PDs, consistent with high cellular turnover and subclonal variation in our NHA E6/E7 model system. Notably, transient peaks in mutation counts emerged at PD 48 (particularly within the C>A and T>C substitutions) which subsequently decreased by PD 73. These data most likely reflect the selectively neutral subclonal expansion of a specific clone, followed by its rapid extinction.

By PD 76, mutational burden increased across all six substitution classes, with the most pronounced gains in C>A, C>T, and T>C mutations (Supplementary Figure 2F). The trinucleotide context of these mutations points to replication-associated processes: C>T gains were enriched at CpG dinucleotides and C>A mutations concentrated at TCA contexts, consistent with clock-like deamination and oxidative damage signatures that accumulate during prolonged culture [25]. Oxidative guanine lesions at telomeric repeats have been shown to drive genomic instability independently of telomere shortening, and the C>A enrichment we observe may reflect a similar oxidative burden genome-wide [26]. This pattern is distinct from the localized APOBEC-mediated kataegis observed at chromatin bridges during telomere crisis [10], and instead reflects the cumulative burden of unrepaired DNA damage in cells dividing through crisis without functional checkpoints, supported by the roughly 60-fold increase in yH2AX expression we observed (Figure S1D).

### Multicolor FISH reveals clonal deletion and subclonal translocation events

To assess the full scope of large-scale structural abnormalities between primary and E6/E7 NHA cells, we employed Multicolor Fluorescence In Situ Hybridization (mFISH) cytogenetic analysis (Figure 3). We generated mFISH karyograms for each bulk and single-cell clone. Primary NHA cells showed no abnormalities across ten unique metaphases (Figure 3A). Conversely, E6/E7 bulk clone #3 showed a consistent deletion of one copy of chromosome 13 in all but one karyogram, supporting the sequencing-based CNV analysis (Figure 3B). We noted numerous complex chromosomal translocations that were invisible to sequencing. We observed t(1q;21q) and t(2q;13q) translocations in respectively 2/10 and 4/10 NHA E6/E7 PD 48 cells. The frequency of t(2q;13q) dropped to only 1/10 at PD 64, whereas a new translocation pattern, t(8q;13q), emerged (Figure 3C).

**Figure 3:**
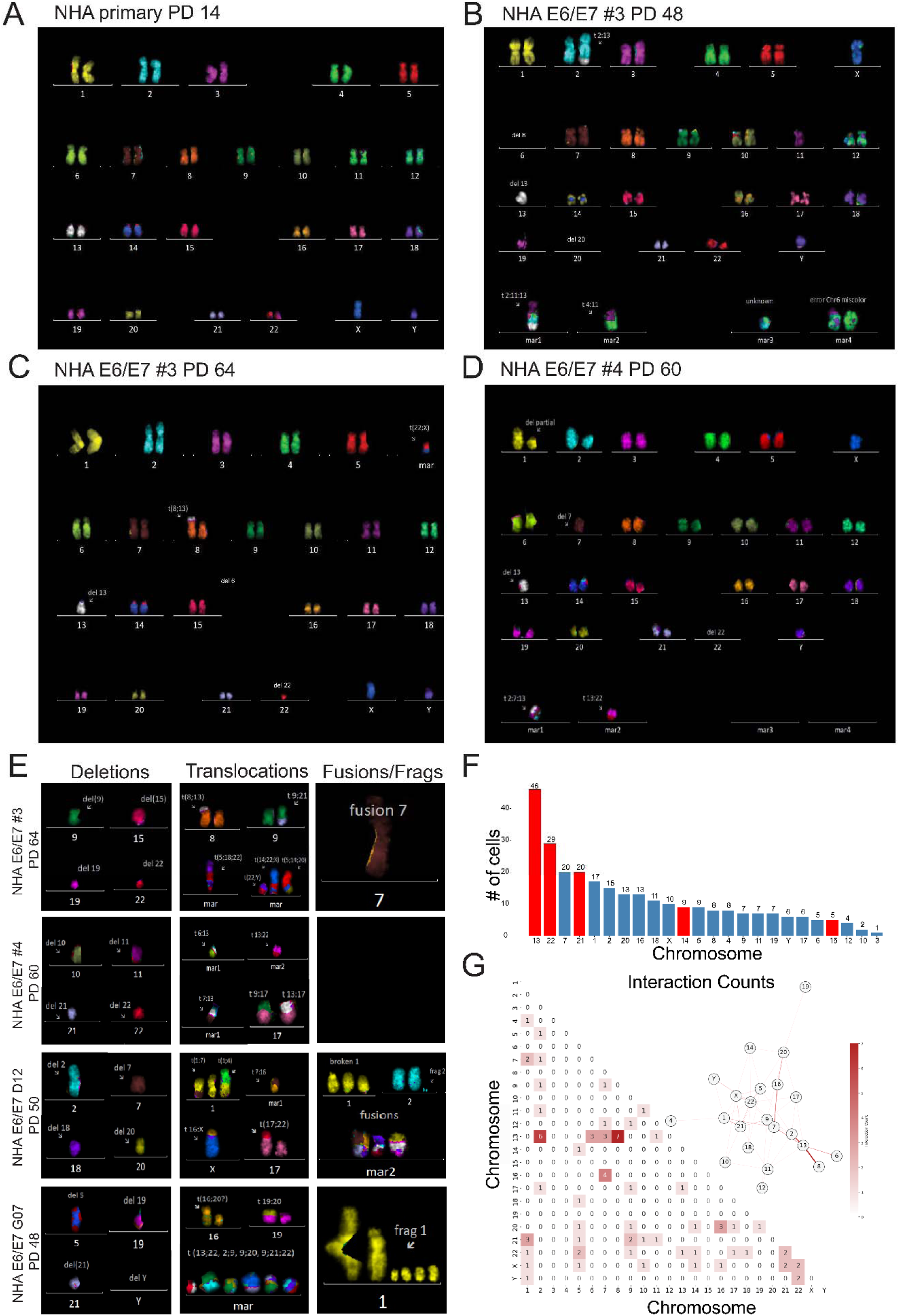
mFISH Karyotype of NHA primary and NHA_E6/E7 metaphase cells at different PDs. **(A)** NHA primary mFISH karyogram showing a standard male karyotype (46, XY). **(B)** Karyogram for NHA E6/E7 bulk clone #3 at PD 48 showing t(2q;13q), del(13), and del(20) along with miscellaneous abnormalities. **(C)** Karyogram for NHA E6/E7 clone #3 at PD 64 showing additional translocations with the appearance of t(8q;13q) **(D)** Karyogram for clone 4 showing chromosome 1, 7, 13, and 22 abnormalities among other features. **(E)** Example abnormalities observed across all NHA E6/E7 bulk and single-cell clones **(F)** Bar chart showing abnormality counts per chromosome across all mFISH scored metaphases. Acrocentric chromosomes are indicated with red bars. **(G)** Inter-chromosomal interaction counts represented as a heatmap and a corresponding network graph. The heatmap displays the frequency of interactions between chromosome pairs, with numerical values and a darker red color gradient indicating higher interaction counts. The inset network visualizes these same relationships; nodes represent individual chromosomes and connecting edges denote interactions, with darker and thicker edges reflecting a greater number of interactions.

NHA E6/E7 clone #4 PD 60 also showed a high percentage of del(13) but demonstrated a completely different spectrum of translocations compared to clone #3 (Figure 3D). Moreover, del(7) was observed in 6/10 #4 PD 60 cells. Clones D12 and G07 often showed fragmentation of chromosomes 1 and 2 but also demonstrated distinct translocations not observed in bulk clones (Figure S3A & B).

Various deletions, translocations and other abnormalities were observed at varying frequencies across each of the bulk and single-cell clones (Figure 3E). Strikingly, acrocentric chromosomes were involved in an unusually high number of these abnormalities, with chromosomes 13, 21, and 22 appearing at higher frequency (Figure 3F). The majority of single-chromosome events were chromosome losses, with partial deletions, fusions and fragmentation events constituting a noticeable minority (Figure S3C). Chromosome 7 abnormalities were biased to single cell clones D12 and G07 (Figure S3D).

To determine whether the enrichment of acrocentric chromosomes among structural abnormalities exceeded expectation, we compared observed abnormality frequencies against a null model weighted by chromosome size. Acrocentric chromosomes (13, 14, 15, 21, and 22) account for 14.6% of the autosomal genome by sequence length yet harbored 41.4% (109/263) of all mFISH-detected abnormalities across 50 scored metaphases, representing a 2.83-fold enrichment (χ² = 151.2, p = 9.5 × 10^-35^). This enrichment was confirmed by a permutation test using a size-weighted null model (100,000 permutations, p < 10^-5^), and was individually significant in each of the five independently derived E6/E7 clones (binomial test, p < 10^-4^ for all five), confirming that the acrocentric vulnerability is a reproducible feature of telomere dysfunction rather than a clone-specific artifact.

Among individual chromosomes, chromosome 22 showed the highest observed-to-expected ratio (6.24-fold), followed by chromosomes 21 (4.68-fold) and 13 (4.40-fold). Notably, chromosomes 14 and 15, while also acrocentric and NOR-bearing, showed no enrichment (O/E = 0.92 and 0.54, respectively), indicating that the vulnerability is non-uniform among acrocentric chromosomes. When restricted to chromosomes 13, 21, and 22 alone, these three chromosomes accounted for 36.1% of all abnormalities despite comprising only 7.4% of the genome (4.90-fold enrichment, binomial test p = 2.0 × 10^-40^). To confirm that this enrichment was not solely driven by the clonal del(13) event, we repeated the analysis after excluding all del(13) events. The acrocentric enrichment remained highly significant (35.0% of events, 2.39-fold enrichment, binomial test p = 5.1 × 10^-15^).

Focusing on inter-chromosomal events and assuming that translocations predominantly occur between neighboring chromosomes, we used our mFISH data to generate a network of chromosome-to-chromosome translocation events (Figure 3G). Chromosomes 13, 7 and 2 exhibited the highest interaction counts, with chromosome 13 forming the most pairwise associations (especially with chromosomes 8 and 2) and maintaining dense connectivity throughout the network, suggesting it occupied a central location in the nucleus and could neighbor many different chromosomes.

### Chromosome-specific FISH characterizes the dynamics of translocations and the persistence of numerical chromosome abnormalities

Because it appeared that translocations in particular were subclonal in our NHA E6/E7 experiment, we sought to more thoroughly assess the temporal dynamics of chromosomal translocations using chromosome-specific FISH probes. Analyzing t(1;21) and t(2;13), which were the most frequent translocations observed in NHA E6/E7 #3 PD 48, we quantified the temporal dynamics of translocations described in Figure 3.

The most striking temporal shift involved t(2;13). Representative metaphase spreads hybridized with chromosome 2 and 13 territory probes illustrated the dramatic change in chr13 translocation partners between PD 48 and PD 64 (Figure 4C). Quantification confirmed that t(2;13) was present in 20/30 (66.7%) of PD 48 metaphases but declined to only 2/30 (6.7%) by PD 64 (Fisher’s exact test, p = 1.9 × 10^-6^; OR = 26.1, 95% CI: 5.0–268.8) (Figure 4D). Metaphase spreads hybridized with chromosome 1 and 21 territory probes similarly illustrated the loss of t(1;21) between time points (Figure 4A). The t(1;21) translocation showed a directionally consistent decline from 4/30 (13.3%) at PD 48 to 0/30 at PD 64, though this difference did not reach statistical significance (Fisher’s exact test, p = 0.11), likely reflecting limited power to detect changes from a low baseline frequency (Figure 4B). The broader burden of translocations involving chromosomes 1 and 21 showed a similar trend, declining from 14/30 (46.7%) to 6/30 (20%; Fisher’s exact test, p = 0.054).

**Figure 4:**
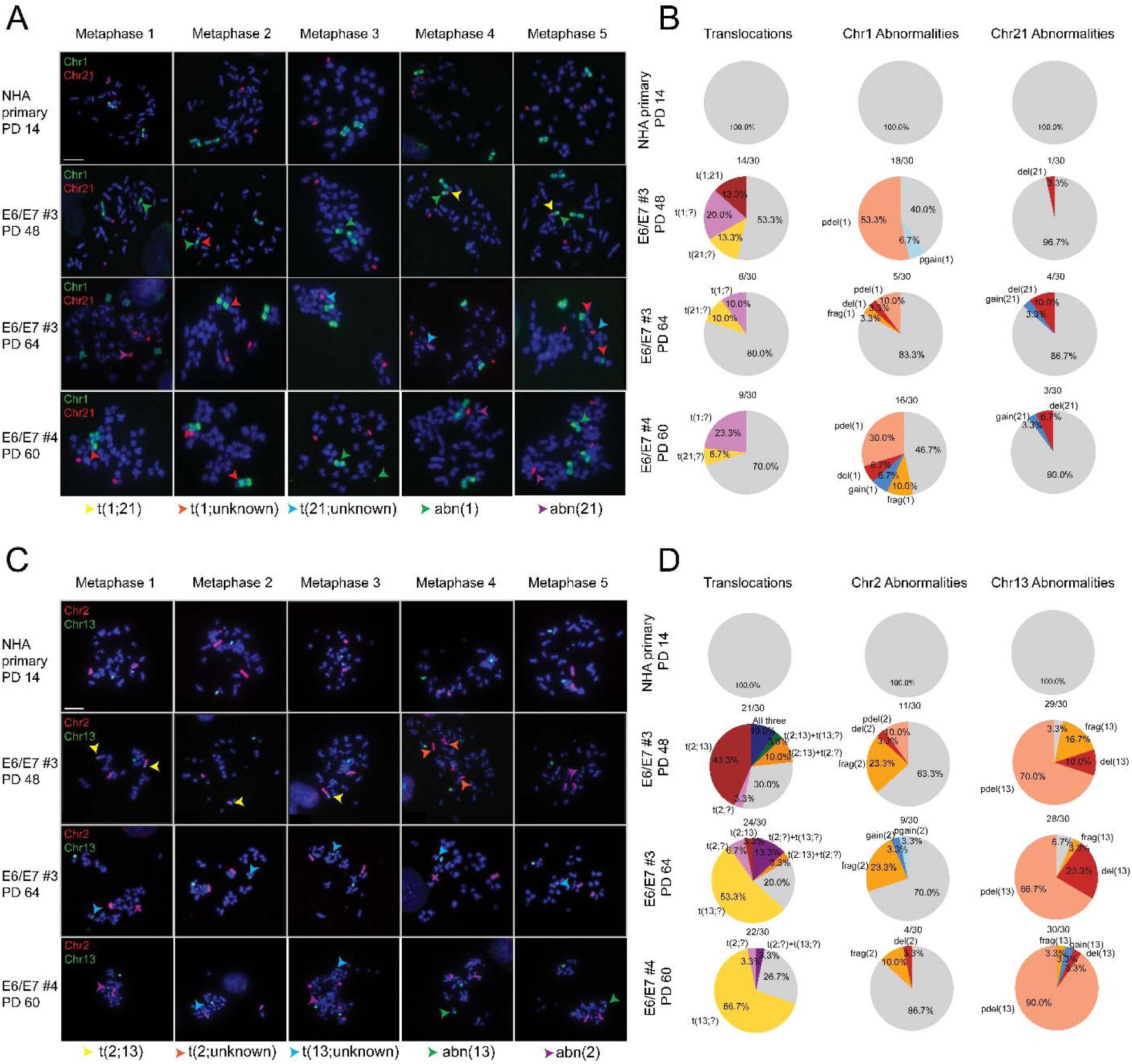
Chromosome Territories for CT1+CT21 and CT2+CT13 in all NHAs. **(A)** & **(C)** Metaphase spreads for NHA primary and each unique clone, hybridized with chromosome territory probes for targeting chromosomes 1 with 21 **(A)** or chromosomes 2 and 13 **(C)** in 5 of 30 cells per clone. NHA primary cells show no abnormalities. Early passage clone 3 (PD 48) shows translocation of chromosomes 2 and 13 at a frequency of 66.7%, indicated by the yellow arrows. The later PD had few cells with t(2;13), but t(8;13) became the predominantly observed phenotype with a frequency of 70%. Clone 4 shows partially deleted chromosome 13, but no distinct pattern of translocation. Scale bar = 10 □m (**B) & (D)** Pie charts showing the frequency of each abnormality in both cases respectively.

In contrast, the overall frequency of chromosome 13 translocation involvement remained stable across all conditions: 21/30 (70%), 24/30 (80%) and 22/30 (73.3%) for #3 PD 48, #3 PD 64 and #4 PD 60, respectively (chi-squared test for heterogeneity, χ² = 0.82, df = 2, p = 0.66). This combination, a highly significant decline in a specific translocation (t(2;13), p < 10^-5^) alongside stable overall chr13 involvement (p = 0.66), identifies chromosome 13 as a preferential substrate for inter-chromosomal rearrangements during telomere dysfunction, whose specific fusion partners are transient.

Other abnormalities involving these chromosomes were more consistent. Deletions showed a contrasting pattern of persistence: del(13) was present in 9/10 mFISH metaphases at PD 48 and 10/10 at PD 64 (Fisher’s exact test, p = 1.0), consistent with clonal fixation. Partial deletions were similarly conserved between time points. Even chromosomal fragmentation events (likely involving micronuclei) showed a stronger persistence between timepoints than translocations, potentially reflecting the reduced fitness imparted by dicentric chromosomes resulting from the latter.

Intriguingly, single-cell clones derived from E6/E7 #3 showed a completely different spectrum of variants. Clone G07 harbored a highly unstable genome with volatile translocation, deletion, gain and fragmentation events (Figure S4A). On the other hand, clone D12 harbored a relatively quiescent genome with a stable deletion of chromosome 13 and fewer other abnormalities (Figure S4B).

Taken together, single-cell cytogenetic analysis revealed a rapidly evolving landscape of subclonal chromosomal abnormalities that are not easily appreciated by genome sequencing. These data argue that traditional cytogenetic techniques such as FISH still hold value in detecting structurally complex and/or subclonal variants that might be missed by conventional short-read sequencing, underlining the need for improved and unbiased approaches for the detection and visualization of genomic abnormalities from sequencing data.

### Telomere dysfunction drives nucleolar reorganization

Noting an unexpected abundance of acrocentric chromosome abnormalities, we sought to establish whether nucleolar organization was similarly disrupted. Because the nucleolus is organized around the nucleolar organizing regions (NORs) that reside on acrocentric short arms, rearrangements of these chromosomes are expected to compromise nucleolar architecture directly. Nucleolar organization also changes with normal replicative aging: small punctate nucleoli are thought to be a hallmark of longevity, whereas enlarged nucleoli are a sign of advanced age or senescence [27–29].

To assess nucleolar integrity, we performed immunofluorescence for the RNA Polymerase I transcription initiation factor UBF, which marks active NORs, and for fibrillarin, the dense fibrillar component (DFC) marker. NHA primary cells displayed 2 to 3 compact, spherical nucleoli per cell with UBF and fibrillarin signal co-localized (Figure 5A-B). Longitudinal quantification further demonstrated a consolidation of nucleolar signal over time: the median number of distinguishable fibrillarin foci decreased from 2 (PD 3) to 1 (PD 27) across primary cells (Kruskal–Wallis, H = 16.42, p = 0.0003), consistent with the nucleolar consolidation characteristic of approaching senescence (Supplementary Figure 5A). In contrast, NHA E6/E7 cells showed a significant increase in nucleolar fragmentation compared to primary cells, with a median of 5 fibrillarin foci (IQR: 3–8) versus 2 (IQR: 1–2) in primary cells (Mann–Whitney U test, W = 1253, p = 3.8 × 10^-16^; Figure 5B). This difference was significant at each matched time point comparison (PD 3 vs. PD 17: p = 1.3 × 10^-3^; PD 14 vs. PD 48: p = 9.2 × 10^-11^; PD 27 vs. PD 60: p = 1.5 × 10^-7^, Supplementary Figure 5C-D). Only 3.3% of primary nuclei (3/90) showed more than four fibrillarin foci, compared to 57.8% of E6/E7 nuclei (52/90; Fisher’s exact test, p = 1.0 × 10^-16^). E6/E7 medians are conservative estimates, as 24.4% of E6/E7 nuclei (22/90) had fibrillarin regions too numerous to count precisely (>8) and were coded as 9 for statistical analysis. Within E6/E7 cells, clone #3 showed a marked increase in fragmentation from PD 17 (median = 3.5) to PD 48 (median ≥ 8.5, with 50% of values censored at >8), while clone #4 at PD 60 (median = 3) was lower than clone #3 PD 48 but still significantly elevated above primary cells. Moreover, nucleoli in NHA E6/E7 cells consistently showed a necklace-like pattern, indicating a breakdown of nucleolar structure. Finally, in a subset of NHA E6/E7 cells we observe a separation of UBF and fibrillarin signal, indicating a decoupling of rDNA processing and transcription and reflecting an even more severe disruption of nucleolar function than structural fragmentation alone. The transition from a small number of consolidated nucleoli to numerous dispersed foci is consistent with a breakdown of the DFC architecture and a failure of NOR clustering.

**Figure 5:**
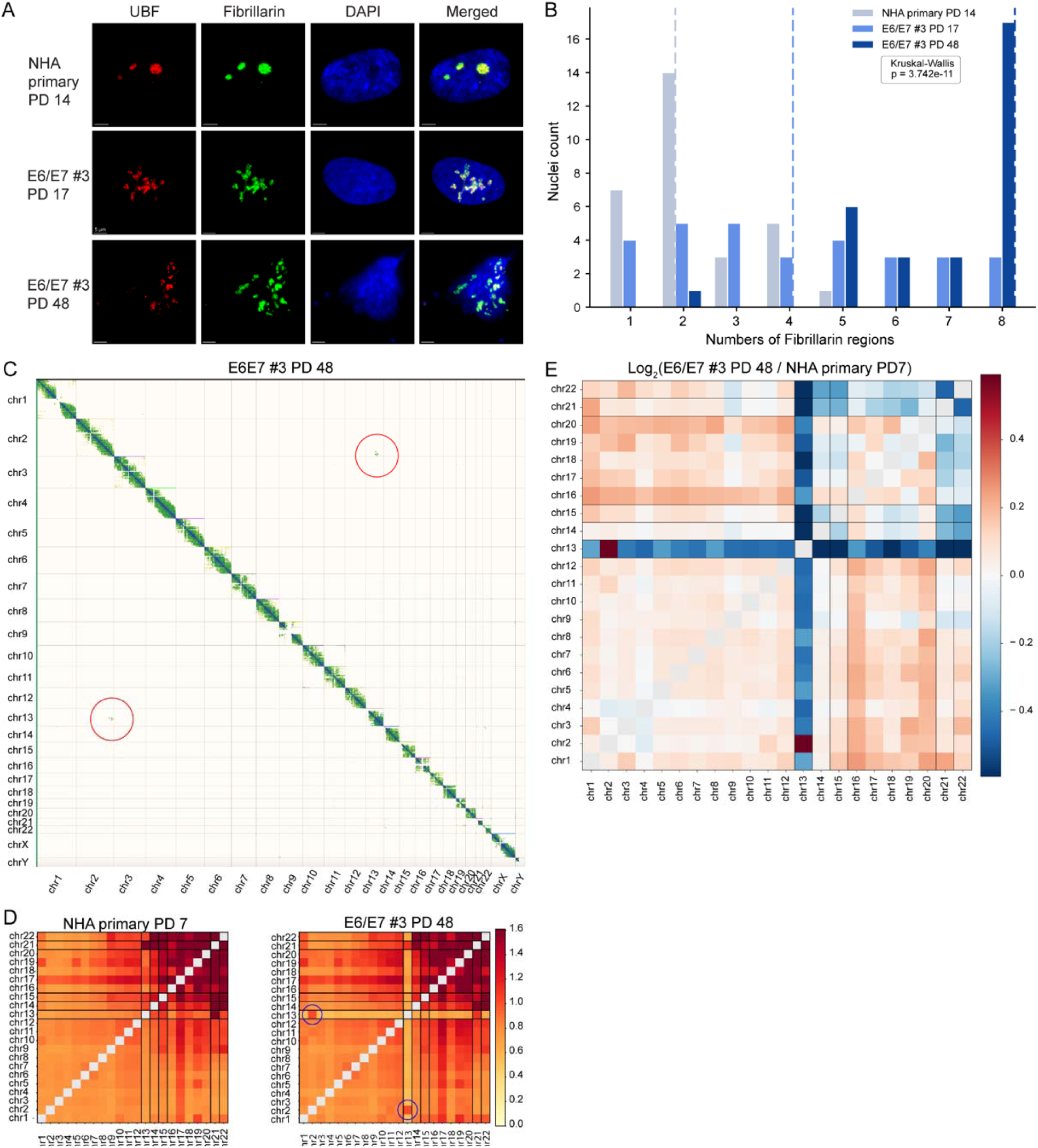
Telomere dysfunction disrupts nucleolar organization in NHA E6/E7 cells. (A) Representative immunofluorescence images of NHA primary cells (PD 14) and NHA E6/E7 #3 cells at PD 17 and PD 48, stained for UBF (red), fibrillarin (green) and DAPI (blue). Merged images shown at right. Scale bar, 5 μm. (B) Bar chart showing the number of fibrillarin foci per nucleus in NHA primary PD 14, NHA E6/E7 #3 PD 17, and NHA E6/E7 #3 PD 48 cells. N = 30 nuclei per condition (90 nuclei total). Kruskal-Wallis p = 3.7 × 10⁻¹¹. (C) Standard alignment-based Hi-C contact map for NHA E6/E7 #3 PD 48. Red circles indicate the only prominent off-diagonal trans-contacts, both corresponding to the chr2:chr13 translocation identified by mFISH (Figure 3B). (D) Inter-chromosomal contact maps generated by KaryoScope for NHA primary PD 7 (left) and NHA E6/E7 #3 PD 48 (right). Values are observed/expected (O/E) normalized contact frequencies. Blue circles highlight chromosome 13 contacts. (E) Log₂FC ratio of O/E-normalized inter-chromosomal contact frequencies between NHA E6/E7 #3 PD 48 and NHA primary PD 7. Blue indicates contact depletion in E6/E7 cells, red indicates enrichment.

The dispersed nucleolar signal in NHA E6/E7 cells suggests a disruption of acrocentric spatial organization. To test this hypothesis, we performed Hi-C on NHA primary and E6/E7 cells across progressive PDs. Traditional Hi-C analysis of this data demonstrated abundant intra-chromosomal interactions and prominent trans-chromosomal contacts involving chromosome 13 representing chromosomal translocations, including the t(2;13) translocation also observed in mFISH (Figure 3B and Supplementary Figure 6).

However, our primary interest was in resolving inter-chromosomal interactions involving the acrocentric chromosomes, that are generally underrepresented in Hi-C and further limited by poor alignments of short reads to the highly repetitive acrocentric short arms. To address this limitation, we applied KaryoScope, a newly developed alignment-free tool from our lab that uses k-mer matching to annotate genomic features including specific chromosomes directly to sequencing reads [30]. Because KaryoScope annotates each read by its k-mer composition rather than by alignment, it retains signal from repetitive regions that mapping-quality and read-support filters in standard pipelines discard. Applying KaryoScope to NHA primary and NHA E6/E7 Hi-C data, we recapitulated the chr2:chr13 translocation also detected by alignment-based approaches (Figure 5D). Importantly, KaryoScope was able to detect substantially larger set of trans-chromosomal interactions, including a cluster of contacts among NOR-bearing acrocentric chromosomes consistent with nucleolar association.

To ask whether NOR contacts are maintained in E6/E7 cells, we computed log₂ fold-changes in inter-chromosomal contact frequency between NHA E6/E7 #3 PD 48 and NHA primary cells (Figure 5E). Contacts among acrocentric chromosomes (13, 14, 15, 21, 22) were markedly reduced in E6/E7 cells, consistent with the disruption of nucleolar organization observed using immunofluorescence. Similar changes in rDNA-containing inter-chromosomal contacts have previously been described as a signature of nuclear reorganization [31]. Chromosome 13 contacts with all other chromosomes were depleted, reflecting the chr13 deletion validated by mFISH, while the chr2-chr13 signal was elevated, consistent with the t(2;13) translocation prevalent at this PD. By PD 76, the chr2-chr13 contact signal had disappeared, consistent with the transient nature of translocations observed by mFISH, while the depletion of acrocentric contacts persisted (Supplementary Figure 6), reflecting ongoing nucleolar disruption.

Together, our cytogenetic (mFISH), immunofluorescence and sequencing data converge on a model in which telomere dysfunction destabilizes acrocentric chromosomes and the nucleolar architecture they collectively form, linking large-scale genomic rearrangements to the disruption of NOR clustering.

## Discussion

The central and most unexpected finding of this study is the disproportionate vulnerability of acrocentric chromosomes during telomere dysfunction. Across 50 scored metaphases from five independently derived E6/E7 clones, acrocentric chromosomes harbored 41.4% of all mFISH-detected abnormalities despite comprising only 14.6% of the autosomal genome, a 2.83-fold enrichment that was highly significant (χ² = 151.2, p = 9.5 × 10^-35^) and individually reproduced in every clone (binomial test, p < 10^-4^ for all five). The enrichment was primarily driven by chromosomes 13, 21, and 22, which together accounted for 36.1% of all abnormalities (4.90-fold enrichment, p = 2.0 × 10^-40^), with chromosome 13 deletions present in over 92% of scored metaphases. Chromosomes 14 and 15, also acrocentric and NOR-bearing, were not enriched. This suggested that acrocentric vulnerability is non-uniform and may depend on factors beyond nucleolar clustering alone, potentially including NOR activity, rDNA copy number, or the intrinsic instability of the rDNA arrays [15]. The most parsimonious explanation for the enrichment of chromosomes 13, 21, and 22 lies in nuclear organization: the NORs on acrocentric short arms cluster within the nucleolus, placing these chromosomes in close spatial proximity during interphase and creating a permissive environment for inter-chromosomal fusions when telomeres become deprotected.

The accompanying transition from compact spherical nucleoli to dispersed necklace-like structures, together with the decoupling of UBF (transcription) and fibrillarin (processing) signals in a subset of E6/E7 cells, links acrocentric rearrangements to a structural breakdown of nucleolar architecture. This phenotype runs opposite to the nucleolar consolidation we observe in primary NHA cells approaching senescence, and the spatial decoupling of UBF and fibrillarin would be expected to impair coordinated ribosome biogenesis. Whether chromosome 13 is preferentially targeted because hemizygous RB1 loss compounds the checkpoint inactivation already achieved by E7, because of differences in NOR activity among acrocentric chromosomes, or a combination of factors, remains an open question. We note, however, that chromosomes 21 and 22, which do not harbor known tumor suppressors with direct relevance to E6/E7, were also strongly enriched (O/E = 4.68 and 6.24, respectively), supporting a model in which spatial proximity within the nucleolus is the primary driver.

To assess whether the morphological changes were accompanied by altered chromosomal organization in the nucleus, we performed Hi-C on NHA primary and E6/E7 cells across progressive PDs. Standard alignment-based Hi-C pipelines exclude reads from rDNA and satellite arrays through mapping-quality and read-support filtering, and therefore underrepresent trans-chromosomal contacts involving the acrocentric short arms. To recover this signal, we deployed KaryoScope, our alignment-free k-mer-based approach described elsewhere [30]. Inter-chromosomal contacts among NOR-bearing acrocentrics were persistently depleted in E6/E7 cells, even as individual translocations such as t(2;13) appeared and disappeared across passages. Together, the contact-based and morphological data support a model in which telomere dysfunction disrupts nucleolar organization at multiple scales: at the protein-component level visualized by immunofluorescence, and at the chromosome-positioning level resolved by Hi-C.

The acrocentric-specific vulnerability we observed was not anticipated when we designed the study, but aligns with prior work demonstrating that acute TRF2 disruption in HCT116 cells produces non-random acrocentric fusions and nucleolar necklace formation, with chromosome 13 as the most commonly involved [18, 19]. Our data extend these observations from acute telomere deprotection in a cancer cell line to chronic, progressive telomere shortening in primary diploid human astrocytes. The convergence across acute TRF2 disruption in transformed cells and chronic telomere shortening in primary astrocytes supports a model in which acrocentric vulnerability is shaped by nucleolar architecture, although broader confirmation across additional cell types and modes of telomere disruption will be needed. Notably, we also observed frequent translocations between acrocentric and non-acrocentric chromosomes (e.g., t(2;13), t(8;13)), which are a class of events largely absent from the acute TRF2 disruption studies, which primarily captured acrocentric-acrocentric fusions [18]. This distinction likely reflects the prolonged window of instability, during which chromosomes beyond the immediate nucleolar neighborhood become exposed to fusion events.

A striking feature of our data is the transient, subclonal nature of many chromosomal translocations. Events such as t(1q;21q) and t(2q;13q) were prevalent at early passages but diminished at later PDs, replaced by entirely new rearrangements. Yet chromosome 13 remained a consistent fusion partner across all conditions, involved in 70–80% of scored metaphases regardless of which other chromosome participated. The persistence of numerical aberrations contrasted sharply with this volatility: deletions and partial deletions were conserved between timepoints, consistent with the reduced fitness imposed by dicentric chromosomes that must segregate at every subsequent division. Single-cell clones derived from the same bulk population displayed entirely different spectra of variants, further underscoring the stochastic character of rearrangements during telomere dysfunction. The decrease in p53 in this model system increases the tolerance for genomic instability, and can enable genome evolution [32, 33].

These observations expose a fundamental limitation of current approaches to studying telomere dysfunction at the genomic level. Mechanistic studies have defined specific pathways of chromatin bridge resolution: nucleolytic processing by TREX1 generating chromothripsis and APOBEC-signature kataegis, and mechanical disruption of the nuclear envelope producing a distinct damage spectrum [9, 10]. These mechanisms predict recognizable genomic signatures detectable by sequencing. Yet longitudinal tracking of structural variants through telomere dysfunction revealed a complexity that exceeds what these mechanisms alone predict [11]. Our mFISH data offer an explanation: the most abundant rearrangements during telomere dysfunction are highly subclonal, transient, and involve large-scale events such as whole-arm translocations, Robertsonian-like fusions and fragmentation events that arise and disappear within a few population doublings. These events are invisible to traditional sequencing approaches because they exist in shifting minor subclones rather than as fixed clonal alterations and require traditional single-cell cytogenetic analysis to be detected. The acrocentric short arms are dominated by satellite and rDNA arrays that are largely inaccessible to conventional bioinformatic methods. Single-molecule long-read sequencing approaches capable of resolving complex rearrangements within these repetitive regions, an emerging direction of our own work, will be critical to fully characterize the genomic consequences of telomere dysfunction.

An important caveat of our experimental design is the use of viral oncoproteins to inactivate p53 and Rb. Beyond checkpoint suppression, E6/E7 exerts pleiotropic effects on the cell and on genome stability: E6 promotes E6AP-dependent degradation of the mitotic kinesin CENP-E, producing polar chromosomes and chromosomal instability independent of telomere dysfunction, and E7 perturbs targets beyond Rb [23, 34, 35]. Some mitotic errors we observe, particularly lagging and polar chromosomes, may therefore reflect direct consequences of oncoprotein expression rather than telomere dysfunction. Interestingly, we do not observe the chromosomal alterations characteristic of HPV-positive squamous cell cancers in our astrocyte model, indicating that cell type likely affects the fitness associated with specific chromosomal alterations [35]. However, the acrocentric-specific enrichment and the nucleolar necklace phenotype are better explained by telomere dysfunction at NOR-bearing chromosomes, as both were also observed following TRF2 disruption in cells expressing neither E6 nor E7 [18, 19]. Future iterations of this model would benefit from CRISPR-based knockout of TP53 and RB1, which would eliminate the pleiotropic effects of the viral oncoproteins and isolate the contribution of telomere dysfunction.

Together, our findings identify the nucleolus as a structural nexus linking telomere dysfunction to large-scale genomic rearrangement. By tracking primary human astrocytes longitudinally from senescence bypass through telomere crisis, we capture an otherwise inaccessible window in which structural damage accumulates and competes under selection. Combined with prior work on acute telomere deprotection in cancer cells [18, 19], our findings indicate that acrocentric vulnerability is a recurrent feature of telomere dysfunction across modes of telomere disruption and cellular contexts. The convergence of cytogenetic, imaging, and chromatin-contact evidence on a single model; points to nucleolar architecture as a determinant of damage patterns and an under-appreciated axis of genomic vulnerability during this instability window. Whether disrupted ribosome biogenesis contributes to the selective pressures of clonal evolution during crisis is the most pressing question raised by this work. More broadly, the link between telomere dysfunction and nucleolar architecture may be relevant to any context in which chromosomally unstable cells expand under altered ribosomal demand, including early tumorigenesis and aging.

## Methods

### Cell culture of Normal Human Astrocytes (NHA)

Normal Human Astrocytes (NHA) and cell culture reagents were obtained from Lonza Bioscience. NHA cells (Cat#CC-2565) were cultured in AGM^TM^ Astrocyte Growth Medium BulletKit ^®^ (Cat. No. CC-3186), passaged using ReagentPack™ Subculture Reagents (Cat. No. CC-5034) and cryopreserved in Cryoprotective Freezing Medium (Cat. No. 12-132A). Cells were maintained in tissue culture treated flasks in humidified incubators set to 37° C, 5% CO2.

### NHA transduction with hTERT viral construct

Early passage NHA cells were transduced with lentivirus made with pLV-hTERT-IRES-hygro, a gift from Tobias Meyer (Addgene plasmid # 85140; http://n2t.net/addgene:85140; RRID: Addgene_85140). Briefly, Lenti-X 293T Cells (Takara Cat. No. 632180) were seeded on a collagen coated (VWR Cat. No. 103700-638) 100mm TC treated dish (Corning Cat. No. 353003). pLV-hTERT-IRES-hygro plasmid DNA was mixed with Lenti-X Packaging Single Shots (VSV-G) (Takara Cat. No. 631275), added to Lenti-X 293T cells, and harvested per manufacturer’s instructions. Virus was used to transduce NHA cells in AGM with 4ug/ml PolyBrene (Sigma-Aldrich Cat. No. T-1003G) overnight. The transduced NHA hTERT cells were then cultured in AGM with 0.3 mg/ml Hygromycin for 6 days to select for cells transduced with the hTERT construct. Aliquots of the NHA hTERT cells were cryopreserved and grown out longitudinally for PD measurements.

### NHA transduction with HPV18 E6/E7 viral construct

HPV18 E6E7 retroviral particles generated in Phoenix amphitropic cells (ATCC Cat. No. CRL-3213) transfected with the plasmid pLXSN18E6E7, a gift from Denise Galloway (Addgene plasmid # 53459; http://n2t.net/addgene:53459; RRID: Addgene_53459) were used to transduce early passage NHA cells. Transduced NHA E6E7 cells were cultured in AGM with 0.8mg/ml Geneticin (G418) for 5 days to ensure presence of E6E7 construct. After 3 days of recovery in AGM, the antibiotic-resistant culture was split into 4 flasks that were then maintained as independent cultures, referred to as Bulk Cultures #1, #2, #3, and #4.

### NHA E6/E7 Single Cell Cloning

After 7 weeks of maintaining the 4 independent cultures, a small number of cells from Bulk Culture #3 were used to perform limited dilution single cell cloning, with a protocol adapted from IDT. Briefly, 100 µL of AGM was added to all wells of a tissue culture treated 96-well plate, except well A1. 200 µL of cell suspension with 4000 cells was added to well A1. 100 µL of cell suspension from well A1 was transferred to B1 and mixed by pipetting. The 1:2 dilutions were repeated down Column 1 using the same pipet tip to complete the first dilution series. 100 µL of AGM was added to wells A–G in Column 1 to reach a final volume of 200 µL/well. Cell suspensions in Column 1 were mixed by pipetting up and down with a multichannel pipettor. The same tips were used to transfer 100 µL of cell suspension across the plate horizontally from Column 1 to Column 2, followed by mixing. The 1:2 dilutions were repeated across the rows using the same pipet tips. 100 µL of conditioned medium (media taken from Bulk #3 after 2-4 days of growth, filtered through a 0.2uM filter) was added to all wells in Columns 1–11 to bring the final volume of each well to 200 µL. This dilution strategy should yield multiple wells that are seeded with a single cell. The plate was incubated in a humidified incubator at 37°C with 5% CO_2_. Wells were monitored with Zeiss Axiovert 40FL microscope to assess growth and determine when cultures were ready to be passaged to larger vessels. Media was replenished weekly, maintaining 50% conditioned media. E6/E7 clones D12, C08, and G07 were grown out longitudinally after this cloning process.

### Longitudinal growth curves

NHA E6/E7 bulk cultures and NHA E6/E7 single cell clones were cultured in T25 and T75 tissue culture treated flasks (VWR Cat. Nos. 82051-074 and 82050-856) in Lonza AGM. Cells were seeded at 5000 cells/cm^2^ and were not allowed to reach confluence. Media was changed every 3-4 days and a Zeiss Axiovert 40FL microscope with 4x and 10x objectives was used to estimate confluency and image cells. When cultures reached 70-90% confluency, the cell monolayers were washed with HEPES Buffered Saline Solution (Lonza Cat. No. CC-5022) and detached with Trypsin/EDTA Solution (Lonza Cat. No. CC-5012) for 3-5 min at 37° C. Trypsin was neutralized by adding an equal volume of Trypsin Neutralizing Solution (Lonza Cat. No. 5002). Harvested live cells were quantified on a Countess™ II FL Automated Cell Counter (Thermo Fisher, Cat. No. AMQAF1000) with trypan blue staining. Cultures were re-seeded and remaining cells were cryopreserved in Lonza cryopreservation media (Lonza Cat. No. 12-132A). At various time points, aliquots of cells were washed with 1x Dulbecco’s Phosphate Buffered Saline (Corning Cat. No. 21-031-CV), collected by centrifugation, and stored as dry pellets at -80° C to be used for downstream assays.

Estimated cumulative population doubling (PD) calculations were performed using:

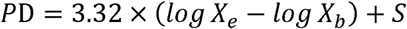

where log is the base 10 logarithm, *X_b_* is the number of live cells seeded at the beginning of the incubation period, *X_e_* is the number of live cells at the end of the incubation period, and *S* is the starting PD (ref ATCC Animal Cell Culture Guide ACCG-042024-v08, p6).

### DNA isolation, PCR-free library preparation, and DNA sequencing

gDNA was isolated from cell pellets using QIAamp DNA Micro Kit (Qiagen Cat. No. 56304) with the addition of 0.5mg/ml RNAse (Qiagen Cat. No. 158922). 500ng of gDNA was sheared on Covaris E220 with parameters designed to target a peak of 300bp (Peak Incident Power (W) = 140, Duty Factor (%) = 10, Cycles per Burst = 200, Treatment time (s) = 80, total volume = 130ul, buffer = Technova Low TE (10mM Tris-HCl, 0.1mM EDTA, pH8.0). ER/AT, adapter ligation, and clean up steps were performed using the Watchmaker DNA Library Prep Kit (Watchmaker Genomics Cat. No. 7K0102-096), IDT for Illumina TruSeq™–Compatible indexed adapters, and AMPure XP Beads (Beckman Coulter Cat. No. A63881). Quality control assays for each library included size measurements by Agilent TapeStation HS D5000 screentape, concentration measurements by Qubit HS dsDNA fluorimetry and KAPA Library Quantification qPCR (Roche Cat. No. 07960336001) prior to pooling. The pool was sequenced on a NovaSeq 6000 (Illumina, Cat. No. 20028312) and NovaSeqX (Illumina, Cat. No. 20104706) and 1% PhiX spike-in to 36X coverage.

### Data processing of sequencing data

Sequencing data was processed using our Snakemake pipeline, available on our GitHub (https://github.com/barthel-lab/general-sWGS). Briefly, empty reads were first removed using Cutadapt (v4.8, minimum length = 1), followed by converting to unaligned BAM format using FastqToSam (GATK4 v4.5.0). Illumina adapters were marked using MarkIlluminaAdapters (GATK4 v4.5.0) and aligned to the human reference genome GRCh38 using BWA-MEM (v0.7.18). PCR duplicates were marked using MarkDuplicates (GATK4 v4.5.0). Aligned reads were filtered for mapping quality ≥ Q30, indexed, and output as BAM files using SAMtools (v1.21).

### Copy number alteration analysis

Copy number analysis was performed in our Snakemake pipeline with the GATK suite (v4.5.0). Processed BAM files were used as inputs. Read counts were collected across 1000 bp intervals using GATK *CollectReadCounts* with the *--interval-merging-rule OVERLAPPING_ONLY* option, using a pre-processed GRCh38 interval list as reference. For sample-to-reference normalization, read counts from NHA E6/E7 clones were denoised against a panel of normals (PoN), which was constructed from primary NHA read counts using GATK *CreateReadCountPanelOfNormals*. The denoising was subsequently performed with GATK *DenoiseReadCounts*, producing both standardized and denoised copy ratio profiles. Allelic counts at heterozygous SNP sites were collected using GATK *CollectAllelicCounts*, with the 1000 Genomes Phase 1 high-confidence SNP site list, obtained via the GATK Resource Bundle from Google Cloud Storage, as the interval input, against the GRCh38 reference genome. Denoised copy ratios and allelic counts were jointly modeled to compute copy number segments using GATK *ModelSegments*. Segmental copy number calls were then assigned using GATK *CallCopyRatioSegments* with default threshold parameters. The copy number profiles across chromosomes were visualized using the Integrative Genomics Viewer (IGV) version 2.19.1.

### Chromosome arm-level aneuploidy analysis

All segmental copy number data from GATK *ModelSegments* for all samples were used for chromosome arm-level aneuploidy analysis. The arm-level copy number alteration calling criterion was adapted from Taylor et al., implemented within our customized R script as part of our Snakemake pipeline [36]. Briefly, the log_2_ copy ratios for all segments were converted into a linear scale. Copy-neutral segments were identified using fixed thresholds (copy ratio between 0.9 and 1.1). A length-weighted mean (*x̅_b_*) and weighted standard deviation (*σ_w_*) were calculated across all copy-neutral segments:

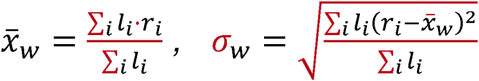

Where *l_i_* is the length, and *r_i_* is the linear copy ratio of segment *i*. Outlier copy-neutral segments were then removed if they fell outside two weighted standard deviations of their call mean:

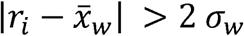

Then, the *̅_x_* and *σ_w_* were recalculated from the remaining segments, yielding the filtered statistics (*x̅_fw_*, *σ_fw_*).

These filtered neutral statistics were then used as z-score thresholds to classify all segments: a segment was called as a deletion (call = −1) if (*r_i_* − *x̅_fw_*) < −2*σ_fw_*, an amplification (call = +1) if (*r_i_* − *x̅_fw_*) > 2 < 2*σ_fw_*, and neutral (call = 0) if the copy ratio fell within the neutral range [0.9, 1.1] or did not exceed the z-score thresholds. All called segments were mapped onto the chromosome arms (GRCh38). Adjacent segments on the same arm sharing the same copy number call were merged into a continuous block. Each arm was then assigned a final copy number call based on the criterion adapted from Taylor et al.: an arm was called a gain (+1) or loss (−1) if the largest contiguous block covered more than 80% of the total arm length (*L_block_* / *L_arm_* > 0.8, otherwise designated as sub-threshold (NA) and excluded from further analysis. The aneuploidy score for each sample was defined as the total number of chromosome arms carrying a definitive gain or loss call. Results were visualized as a heatmap with samples hierarchically clustered based on Euclidean distances computed from the arm-call matrix, using Ward’s minimum variance method (ward.D2).

### Analysis of somatic single nucleotide variant and indels

Somatic SNVs and indels were identified using GATK *Mutect2* in tumor-normal mode. Briefly, processed BAM files were analyzed with NHA E6/E7 clones assigned as the “tumor” (query) and the parental NHA cell line assigned as the “normal” for filtering pre-existing background variants. F1R2 read counts were collected during the *Mutect2* calling step and used to train a read orientation model via GATK *LearnReadOrientationModel*. This model was subsequently applied during variant filtering with GATK *FilterMutectCalls*, and only variants passing all filters (PASS) were retained and exported in VCF format. The distributions of variant allele fraction for each E6/E7 clone were visualized as ridgeline plots using matplotlib version 3.10.0 in Python version 3.12.8. The sharing and uniqueness of variants across serial passages within each clone were visualized as UpSet plots using the UpSetR package in R.

### Alignment-based Hi-C data processing and analysis

Hi-C reads were aligned to the T2T-CHM13v2.0 reference and processed according to the Arima Mapping pipeline (A160156 v03, https://github.com/ArimaGenomics/mapping_pipeline). In summary, Hi-C reads were aligned using BWA mem with default parameters (version 0.7.18-r1243-dirty). The bam files were filtered using the filter_five_end.pl script to retain only the portion of chimeric reads that map in the 5’-orientation. The two_read_bam_combiner.pl script was used to combine the single-end Hi-C reads. Picard (version 3.1.1, https://github.com/broadinstitute/picard) was used to add read groups to the bam files and remove optical duplicates. Hi-C contact maps were created with PretextMap (version 0.1.9, https://github.com/sanger-tol/PretextMap) and visualized with PretextView (version 0.2.5, https://github.com/sanger-tol/PretextView).

### Hi-C inter-chromosomal contact analysis with KaryoScope

In addition to standard alignment-based Hi-C pipelines, we applied KaryoScope an alignment-free k-mer matching method that annotates each read by its k-mer content [30]. Per-chromosome KMC databases were constructed from the CHM13 v2.0 reference assembly, with one database per chromosome (chr1–22, chrX, chrY). For each Hi-C read, KaryoScope matched constituent k-mers against the databases using the *get_featureIDs* utility, and adjacent k-mer hits were consolidated into chromosome intervals along the read by the smoothing step, producing a BED file of chromosome assignments per read. Per-read chromosome pair counts were then aggregated across all reads in a sample to yield a pairwise chromosome co-occurrence table. KaryoScope is available at https://github.com/barthel-lab/KaryoScope.

### Contact matrix construction

For each sample, the co-occurrence table was assembled into a symmetric chromosome × chromosome count matrix *M* restricted to chr1–chr22, with each unordered pair contributing its count to both *M_A,B_* and *M_B,A_*. The diagonal was then masked to exclude intra-chromosomal contacts, so that all subsequent normalization steps operate on inter-chromosomal contacts only.

### Size-normalized contact density

To compare inter-chromosomal contact frequencies across chromosome pairs and across samples, we computed a per-pair density that normalizes each pair count by the product of the two chromosome lengths and by the sample’s total inter-chromosomal pair count. This removes both the size-product bias inherent in inter-chromosomal contact frequency, in which larger chromosomes accumulate more contacts purely by virtue of their physical extent, and library-composition differences between samples:

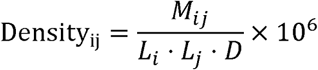

where *L_i_* and *L_j_* are the CHM13 v2.0 chromosome lengths in Mb, and *D* is the total inter-chromosomal pair count in the sample, computed from the count matrix as 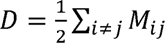 to correct for the symmetric double-counting in *M*. We restricted the denominator to the data pool used in the analysis (chromosome-annotated inter-chromosomal pairs) rather than the full library so that samples are compared on the basis of their in-scope reads only.

### Observed / Expected normalization

To place chromosome pairs within a sample on a common scale, each density value was divided by the mean of the sample’s off-diagonal density values:

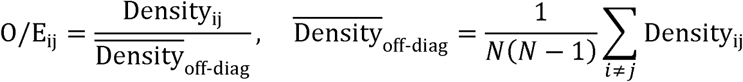

where *N* = 22 is the number of chromosomes in the matrix. The expected value is therefore the typical inter-chromosomal contact density for that sample, computed from the off-diagonal pairs only. An 0/E_ij_ value of 1.0 corresponds to a chromosome pair at the typical inter-chromosomal contact density, values greater than 1.0 to chromosomes pairing more frequently than expected, and values less than 1.0 to chromosomes pairing less frequently than expected.

### Differential contact analysis

For each E6/E7 sample, log_2_ fold-changes in O/E versus NHA primary cells were computed pair-wise as

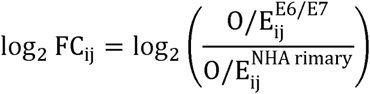

Library depth and chromosome-size effects cancel in this ratio, leaving signal that reflects specific changes in inter-chromosomal contact preference. Chromosome pairs were filtered when both samples accumulated fewer than 10 reads supporting that pair. Per-clone log_2_ fold-change matrices were computed for each E6/E7 sample.

### Heatmap rendering

O/E matrices (Figure 5D and Supplementary Figure 7A) were displayed as 22 × 22 heatmaps spanning chr1–chr22 using a sequential YlOrRd colormap. Per-clone log_2_ fold-change matrices (Figure 5E and Supplementary Figure 7B) were displayed using a diverging RdBu colormap centered at zero (red for enrichment in E6/E7, blue for depletion). In both cases, a shared color-scale upper limit was applied across all panels of the same type so that any two panels of that type can be compared directly. The diagonal was masked from display, the y-axis was inverted so that chr22 appears at the top, and acrocentric chromosomes (chr13, chr14, chr15, chr21, chr22) were outlined as rows and columns to guide the eye to NOR-bearing chromosomes.

### TRF Analysis

Telomere length was determined by telomere restriction fragment (TRF) analysis as previously described [37]. Genomic DNA was extracted with QIAamp DNA Micro Kit (Qiagen Cat. No. 56304) with the addition of 0.5mg/ml RNAse (Qiagen Cat. No. 158922 and quantified using a Qubit Fluorometric Quantification (Qubit 3.0 Fluorometer, Thermo Fisher Scientific). For each sample, 600 ng of genomic DNA was digested overnight (∼12 h) at 37°C with HinfI (New England Biolabs, R0155S) and RsaI (New England Biolabs, R0167L). The resulting DNA fragments were resolved on a 0.7% agarose gel in 1× TAE buffer.Electrophoresis of constant-field gel was performed for 16 h at 2 V/cm in the cold room.Following electrophoresis, the gel was dried for 2 h at 42°C and stained with EthidiumBromide (Sigma-Aldrich, 2375) and imaged with BioRad GelDoc XR imager (BioRad).The gel was then subjected to sequential in-gel processing: denaturation for 30 min in 0.5 M NaOH, 1.5 M NaCl, followed by neutralization for 30 min in 0.5 M Tris-HCl (pH8.0), 1.5 M NaCl. Subsequently, the gel was prehybridized with 10x Denharts’ Buffer, 5xSSC and 0.5% SDS, then hybridized overnight at 42°C with a telomeric C-rich probe 2. The hybridized gel was then washed 3 times with 2× SSC and 0.5% SDS for 15 mins each and 3 times with 2× SSC + 0.1% SDS for 15 mins each. Finally, the gel was exposed to a PhosphorImager Screen (GE Healthcare) and imaged on a Typhoon scanner (GE healthcare) ImageQuant (Molecular Dynamics). Telomere length was analyzed using ImageQuant.

### Protein lysate preparation

Cell pellets of 0.6 - 2.0 x 10^6^ cells were harvested in 1.5ml microfuge tubes at 300xg for 5 min, followed by a single wash of 1x PBS. Cells were resuspended in 400 ul of 1x RIPA Buffer (Cell Signaling Technology Cat. No. 9806) with 1x complete Protease Inhibitor (Sigma-Aldrich Cat. No.4693116001). Samples were sonicated for 10 seconds at 50% amplitude on ice, followed by centrifugation at 15,000xg for 10min at 4°C to spin down cell debris. One aliquot of cleared lysate was used to quantify protein concentration with BCA assay (Thermo Fisher Cat. No. A55865). A second aliquot of 200ul of cleared lysate was transferred to 1.5ml Protein LoBind tubes (Eppendorf Cat. No. 022431081) containing 100ul of 3x Blue Loading Buffer (Cell Signaling Technology Cat. No. 56036) with 3x DTT (Cell Signaling Technology Cat. No. 14265). The samples were heated to 95C for 10min to denature and were stored at -80°C.

### Western blot

Samples were run on 12% acrylamide gels, transferred to nitrocellulose overnight, and blocked in 5% milk in TBS + 0.1% Tween 20 (TBST) for 1 h at room temperature before overnight incubation with primary antibodies, which were diluted in 2% bovine serum albumin + 0.02% sodium azide in PBS. Primary antibody dilutions: p53 1:500 (DO-1; Cell Signaling Technology), P-Histone H2A.X 1:1000 (20E3; Cell Signaling Technology), and vinculin 1:2000 (hVIN-1; Milliporesigma). Blots were washed 3 times in TBST, then incubated for 45 m in secondary antibodies which were diluted 1:10,000 (Licor IRDye) in 5% milk in TBST. Blots were imaged on a LI-COR ODYSSEY XF Imager and quantified with ImageStudio.

### Multicolor Metaphase Fluorescence In Situ Hybridization (mFISH)

NHA primary and E6/E7 clones for karyotyping were prepared following manufacturers’ recommendations. Briefly, cells were cultured to 70% confluency and treated with 0.2ug Colcemid to arrest nuclei in metaphase. Cells were harvested with 0.5% Trypsin-EDTA (Gibco, Cat. No. 15400054), collected by centrifugation at 1,500 rpm for 5 minutes, and resuspended with 0.38% KCl (Invitrogen, Cat. No. AM9640G) for 20 minutes. Swollen cells were resuspended in Ice-cold Methanol: Acetic Acid fixative and dropped on humidified slides. Slides were incubated for 30 minutes in 2X SSC warmed to 70℃. Concurrently, 10µL of 24XCyte human multicolor probe mix (MetaSystems, ref D-0125-060-DI) were denatured at 75℃ for 5 minutes followed by incubation for 30 minutes at 37℃. Slides were removed from the warmed 2X SSC solution and slowly cooled to room temperature. The slides were then incubated in room 0.1X SSC, fresh 0.07N NaOH, 0.1X SSC again, 2X SSC, and dehydrated with an ethanol series. 10µL of the probe mixture was then applied to each slide and allowed to incubate at 37℃ for 48 hours. Post hybridization washes were carried out according to manufacturer’s recommendations. Slides were mounted using DAPI antifade (MetaSystems, ref D-0902-500-DA) and covered with 22×22 #1.5 glass coverslips. Images were obtained using a Zeiss AxioImager Z2 equipped with the MetaSystems *Metafer5* Slide Scanning Platform software (MetaSystems, *Metafer5*).

### Karyogram Generation

Karyograms were generated with the Interactive Karyotyping system (Ikaros) module, a feature of the *Metafer5* system. Captured images were processed by background subtraction, object thresholding, object separation, and pixel calling. False colors were assigned to each chromosome for clear identification. Chromosome assignments were confirmed by comparing fluorescent spectra to the manufacturer key and each chromosome was matched to its respective ideogram. Ten PDF reports for each cell line were generated and exported from the software.

### Statistical Analysis of Acrocentric Enrichment

To test whether acrocentric chromosomes were disproportionately affected by structural abnormalities, we compared the observed frequency of abnormalities on acrocentric chromosomes (13, 14, 15, 21, 22) versus non-acrocentric chromosomes against a null distribution weighted by chromosome size (GRCh38 assembly lengths). Each autosome was counted at most once per metaphase (presence/absence of any abnormality). A chi-squared goodness-of-fit test was performed against size-weighted expected proportions. Additionally, a one-sided binomial exact test was performed under the null hypothesis that the proportion of acrocentric abnormalities equals the fraction of the autosomal genome occupied by acrocentric chromosomes (14.6%). A permutation test was conducted with 100,000 iterations, in which each observed abnormality was randomly assigned to a chromosome with probability proportional to its sequence length, and the number of permutations yielding an equal or greater number of acrocentric abnormalities was used to calculate the empirical p-value. To assess reproducibility, the binomial test was repeated independently for each of the five E6/E7 clones. A sensitivity analysis was performed by excluding all del(13) events to confirm that the enrichment was not solely driven by the clonal chromosome 13 deletion. Analysis was performed in R (v4.5.0) using base statistical functions.

### Statistical Analysis of Translocation Dynamics

The temporal dynamics of specific translocations were assessed using two-sided Fisher’s exact tests on 2×2 contingency tables comparing the frequency of each translocation event between PD 48 and PD 64 in clone #3 (n = 30 metaphases per time point). The stability of overall chromosome 13 translocation involvement across three independent conditions (clone #3 PD 48, clone #3 PD 64, and clone #4 PD 60) was assessed using a chi-squared test for heterogeneity on a 3×2 contingency table. Analysis was performed in R (v4.5.0).

### Chromosome Territory PAINT FISH

Premixed whole chromosome PAINT probes from MetaSystems for chromosome 1, 2, 13, and 21 (MetaSystems, Cat#D-03001-100-FI, D-0302-100-OR, D-0313-100-FI and D-0321-100-OR respectively) were hybridized to metaphase spreads based on the manufacturer’s protocol. Thirty metaphases were imaged per cell line. Images were captured at 60X magnification using a Nikon Eclipse Ti2-E widefield microscope. Analysis was performed by counting the number of translocations, deletions, and other abnormalities seen in E6/E7 cells compared to primary NHA.

### Immunofluorescence microscopy

For quantification of mitotic errors, cells (normal NHA and E6/E7 bulk clones #3, 4, and single cell clones G07 and D12) were grown on glass coverslips and fixed in 4% PFA when confluent, and stained for α-tubulin and DAPI as described in [38]. Images were taken Nikon Eclipse Ti2-E widefield microscope with a 100x/1.4 numerical aperture oil objective. Images are maximum projections of deconvolved 0.2 μm z-stacks.

For nucleolar analysis, normal NHA cells, E6/E7 bulk clones from batch #3, and bulk clones from batch #4 were seeded in chamber slides. Cells were washed with 1X PBS, fixed with 4% paraformaldehyde (PFA) for 10 minutes and permeabilized with 0.5% Triton-X100 for 5 minutes. Cells were then blocked with UltraCruz Blocking Reagent (SantaCruz Technologies, cat# sc-516214) for 30 minutes. Primary antibodies were incubated for 30 minutes at 37C (rabbit anti-Fibrillarin 1:1000 and mouse anti-UBF 1:150) then washed 2x with PBS. Secondary antibodies were also incubated for 30 minutes at 37C (AlexaFluor goat anti-rabbit 488 and goat anti-mouse 555, at 1:800). Cells were washed 1x with PBST, 2x with PBS, dehydrated with an ethanol series, and finally mounted with DAPI antifade (MetaSystems, cat#D-0902-500-DA). Thirty interphase images per cell line were captured at 60X using a Nikon Eclipse Ti2-E widefield microscope.

Distinguishable fibrillarin foci were counted in each of the 30 interphase nuclei per condition. In E6/E7 cells with highly fragmented nucleoli, foci too numerous to count precisely were recorded as “>8” and conservatively coded as 9 for statistical analysis; this approach biases against the hypothesis of increased fragmentation, ensuring that any significant result is robust. Group comparisons of fibrillarin foci counts were performed using two-sided Mann–Whitney U tests; within-group trends across PDs were assessed by Kruskal–Wallis tests. The fraction of nuclei exceeding four fibrillarin foci was compared between groups using Fisher’s exact test.

## Author Contributions

O.M. performed chromosome territory FISH, nucleolar immunofluorescence, karyotyping, and figure generation. Y.C. and Y.H. analyzed the data and generated figures. N.F. performed cell culture and generated growth curves. Y.H., Y.C., R.R.B., M.K. conducted short-read sequencing analysis. P.F.C., N.M. and A.B. conducted mitotic error immunofluorescence and performed analysis. S.C and T.Z. performed TRF analysis. P.C., T.Z., and F.P.B provided advice and direction for experiments. O.M., Y.C., Y.H. and F.P.B wrote the manuscript. F.P.B. conceived and supervised the project. All authors reviewed, commented on and approved the final paper.

## Acknowledgements

This work was supported by the Department of Defense (award HT9425-23-1-0844 to F.P.B.), the NIH (K08CA256166 to P.F.C and R35GM162212 to T.Z.), and CEP award from the Head and Neck Cancer SPORE (P50CA278595 to P.F.C), and Radiological Society of North America (Research Scholar Grant to P.F.C.). Large language models were used in the preparation of this manuscript to improve grammatical accuracy and readability. The authors reviewed all edits and retain sole responsibility for the accuracy and integrity of the content. This research includes work performed in TGen’s Collaborative Sequencing Center; a City of Hope Comprehensive Cancer Center supported shared resource (NCI-P30CA033572). The authors thank Thomas Dennis and Jeff Sanford for helpful discussions and assistance with experimental optimization.

## Supplementary Figures

**Supplemental Figure 1:**
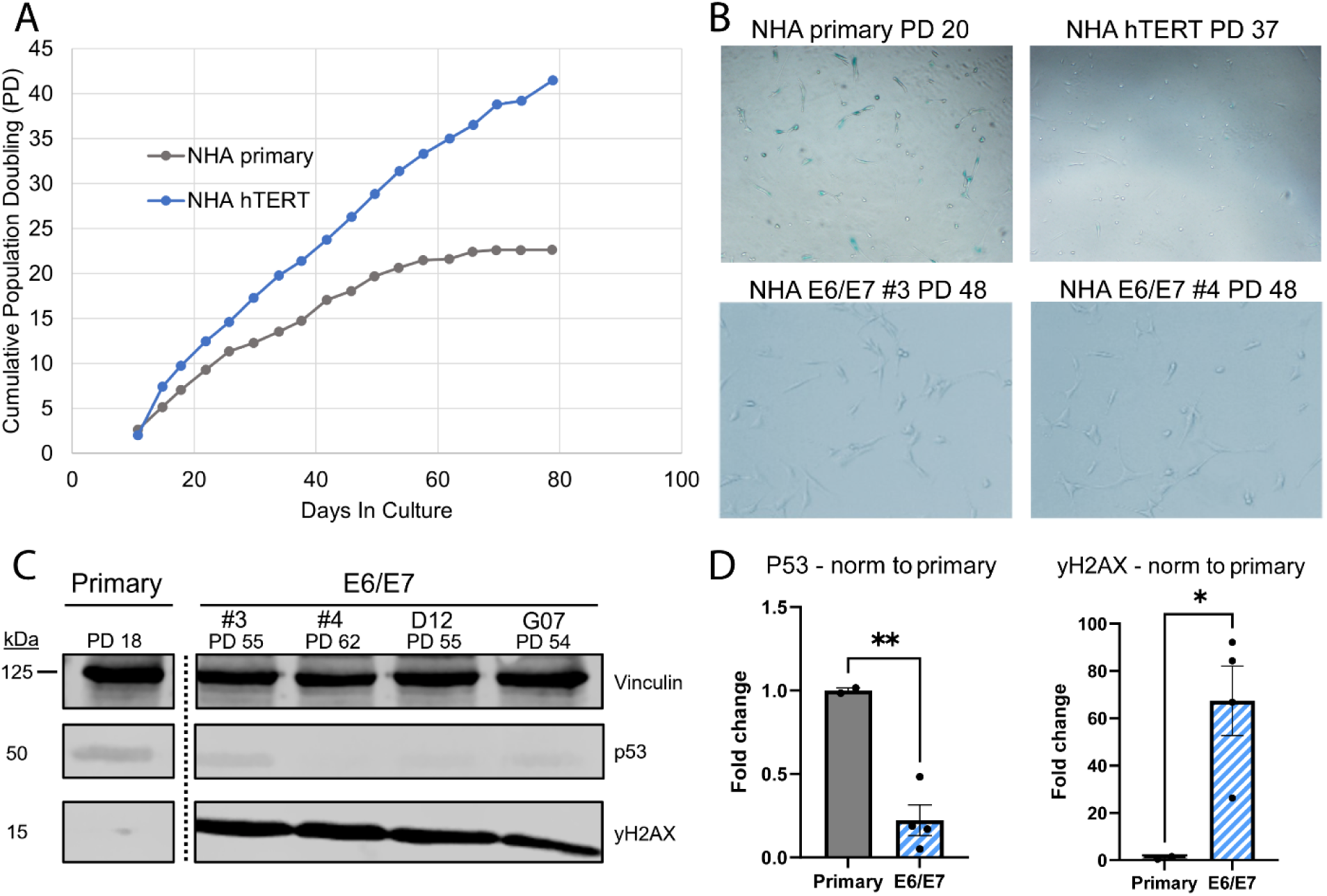
E6/E7 model survives past primary senescence points while DNA damage markers increase significantly. **(A)** Growth curves of NHA primary vs NHA hTERT. The population doubling of the primary line plateaus at PD22, while hTERT continues to grow past PD 40 at day 80. **(B)** Beta-galactosidase staining of NHA primary, hTERT, clone #3, and #4 to assess senescence. Primary NHA cells began to senesce at PD 20 as indicated by blue staining within individual cells, while NHA hTERT and the E6/E7 transduced clones show no signs of senescence **(C)** Western blot of NHA primary vs E6/E7 measuring p53 levels and yH2AX levels at late passages. **(D)** Quantification of western blot data confirms significantly low levels of p53 and high levels of yH2AX in primary vs E6/E7.

**Supplemental Figure 2:**
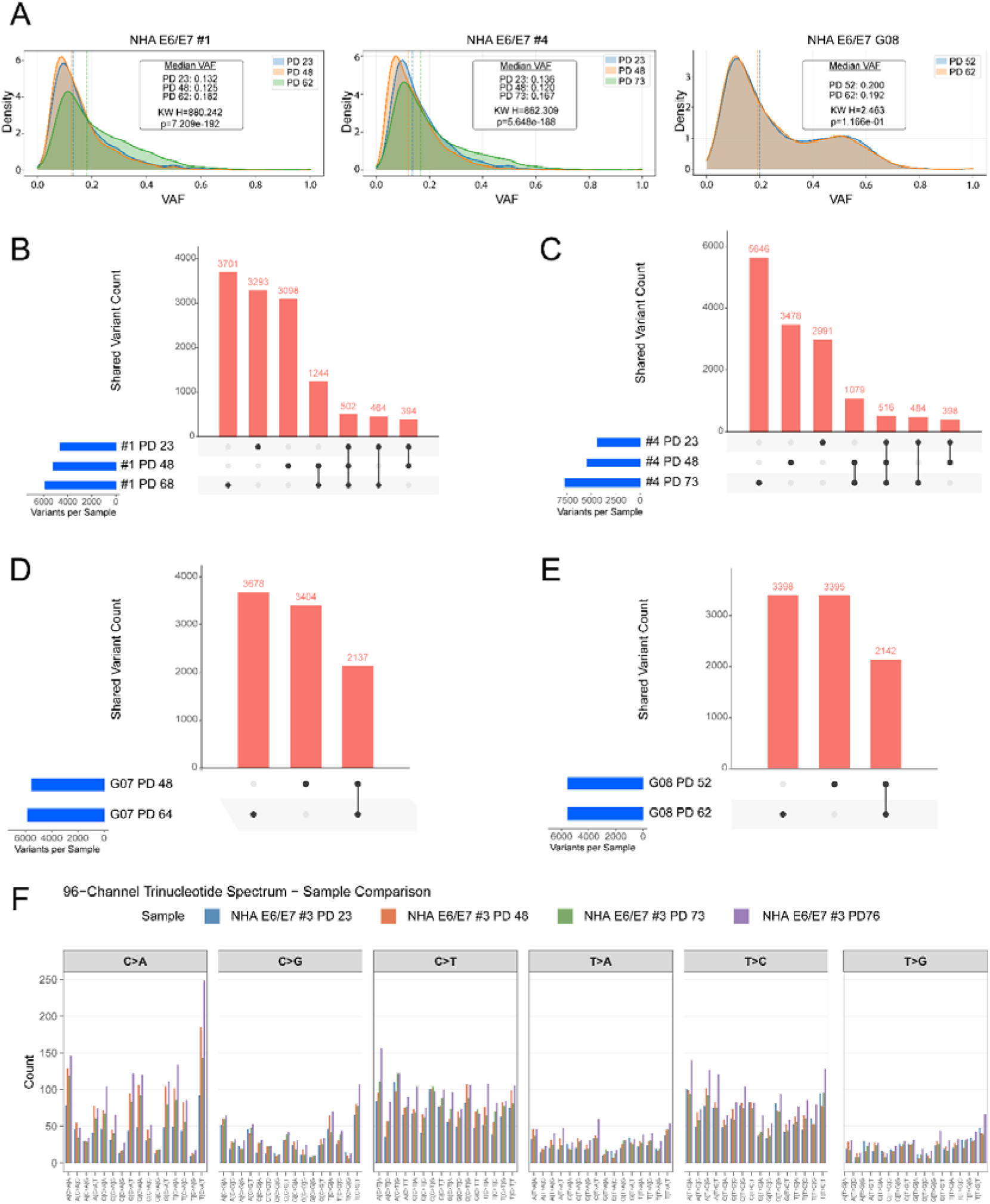
Additional VAF distribution data and upset plots for remaining clones. **(A)** Variant allele frequency (VAF) density distributions for representative samples (bulk #1, #4, and single-cell lone G08) at progressive PDs. Dotted vertical lines denote median VAFs, with Kruskal-Wallis (KW) test statistics provided for each clone. (B-E) UpSet plots illustrating the intersection of shared and unique SNVs and indels across multiple PDs for bulk clone #1 **(B)**, #3 **(C)**, and single-cell clones G07 **(D)** and G08 **(E)**. **(F)** The 96-channel single-nucleotide substitution (SNS) spectrum is shown for bulk clone #3 at PD 23, 48, 73, and 76. Mutations are grouped by substitution type: C>A, C>G, C>T, T>A, T>C, and T>G, and further subdivided by 5’ and 3’ flanking nucleotides. The Y-axis indicates raw mutation count. Colors denote individual passage time points.

**Supplemental Figure 3:**
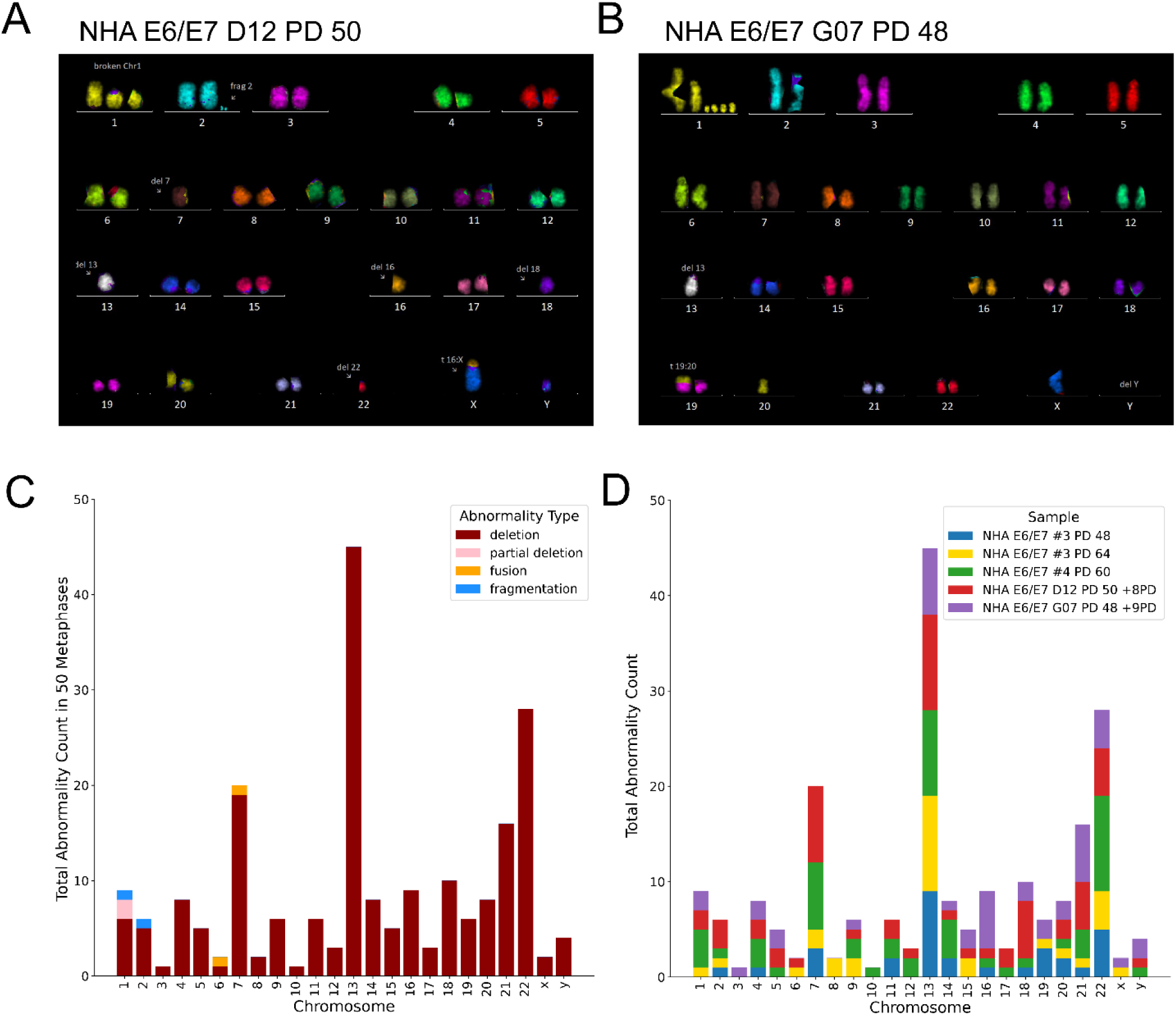
24-color mFISH for additional NHA E6/E7 clones. **(A)** Clone D12 shows chromosome 7,13, 16, 18, and 22 deletion, and translocation of chromosome 16 to chromosome X. (**B)** Clone G07 showing fragmentation of chromosome 1, deletion of chromosome 13, 20, & Y, and translocation of chromosomes 19 with a residual segment of 20. (**C)** Abnormalities per chromosome broken down by four major types. Deletions were predominantly seen across all chromosomes. Partial deletions, fusions, and fragmentations occurred at roughly equal counts (**D)** Count of abnormalities seen within each chromosome across specific clones. Abnormality counts were not equally proportional in all samples.

**Supplemental Figure 4:**
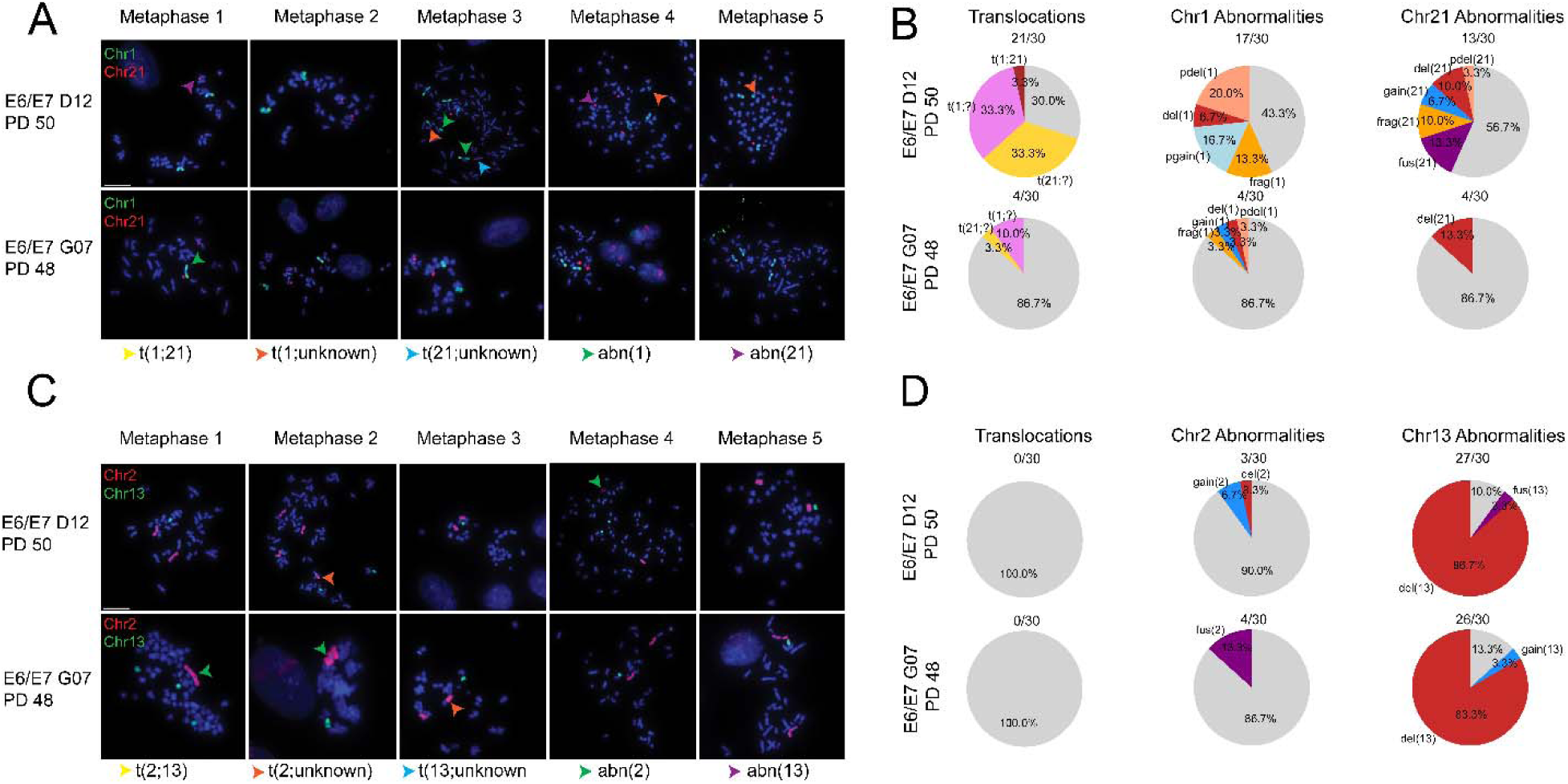
Additional CT images for D12 and G07. **(A)** Clone D12 and G07 metaphases hybridized with CT1 and CT21. Arrows denote abnormality types seen within each cell, with yellow arrows marking t(1;21), orange marking t(1;unknown), blue marking t(21;unknown), green marking chromosome 1 abnormalities, and purple marking chromosome 21 abnormalities (**B)** Analysis of translocations and abnormalities observed in chromosome 1 and 21. Translocations involving either chromosome are noted. Those involving chromosomes 1 or 21 and unknown chromosomes were seen at 33.3% each. Those involving chromosomes 1 and 21 specifically were only seen in 3.3% of cells (n=30). **(C)** Clone D12 and G07 metaphases hybridized with CT2 and CT13. Arrows denote abnormality types seen within each cell, with yellow arrows marking t(2;13), orange marking t(2;unknown), blue marking t(13;unknown), green marking chromosome 2 abnormalities, and purple marking chromosome 13 abnormalities **(D)** analysis of translocations and abnormalities observed in chromosome 2 and 13. No translocations were seen for either chromosome, but deletion of chromosome 13 became prevalent. Fusions appeared in both single cells clones at low rates.

**Supplementary Figure 5:**
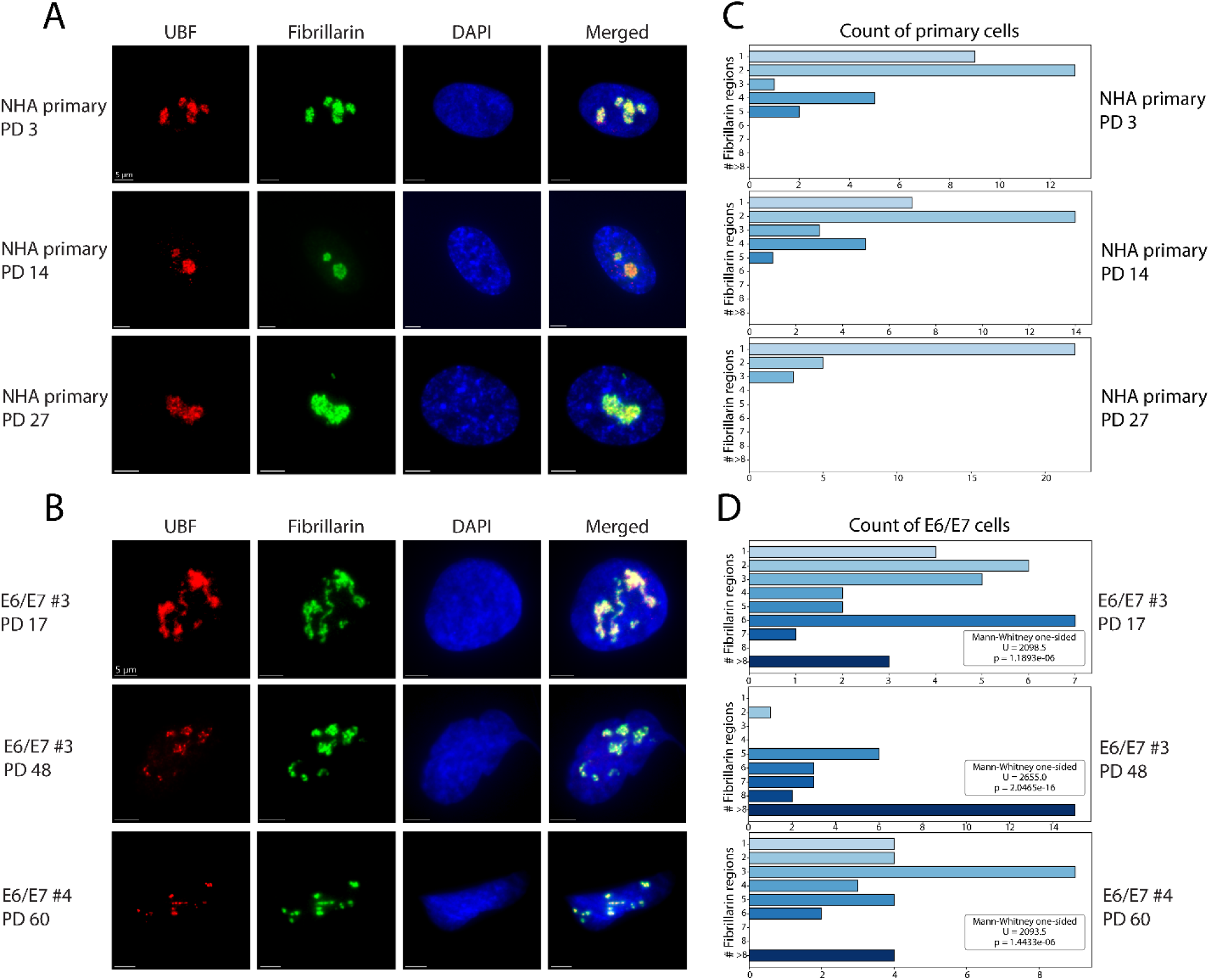
Telomere dysfunction disrupts nucleolar organization in NHA E6/E7 cells. **(A)** NHA primary cells immunostained with fibrillarin (green) and UBF (red) antibodies at three different PDs representing young, middle, and old timepoints (PD 3, 14, and 27 respectively). Primary cells show distinct spherical boundaries denoting the nucleoli **(B)** E6/E7 cells were similarly stained with fibrillarin and UBF at different timepoints (PD 17, 48, and 60). The spherical boundaries unraveled into more linear structures that covered a wider area of the nucleus. **(C)** Number of distinguishable fibrillarin regions in NHA primary cells at each time point n=30 per time point. **(D)** Number of distinguishable fibrillarin regions in E6/E7 cells at each time point n=30 per time point.

**Supplementary Figure 6:**
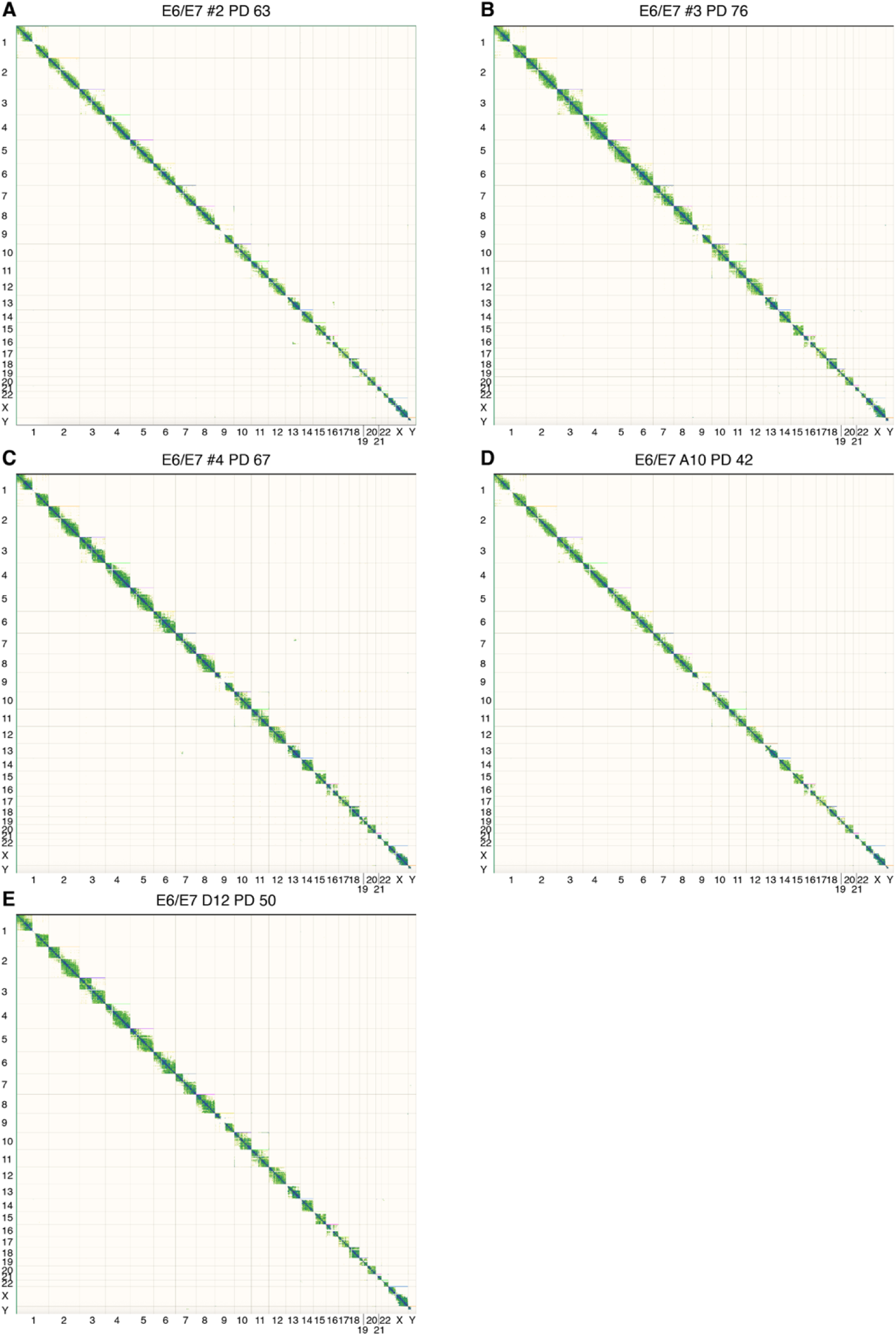
Standard Hi-C contact maps for additional NHA E6/E7 samples. Genome-wide alignment-based Hi-C contact maps for NHA E6/E7 #2 PD 63 (A), NHA E6/E7 #3 PD 76 (B), NHA E6/E7 #4 PD 67 (C), NHA E6/E7 A10 PD 42 (D) and NHA E6/E7 D12 PD 50 (E).

**Supplementary Figure 7:**
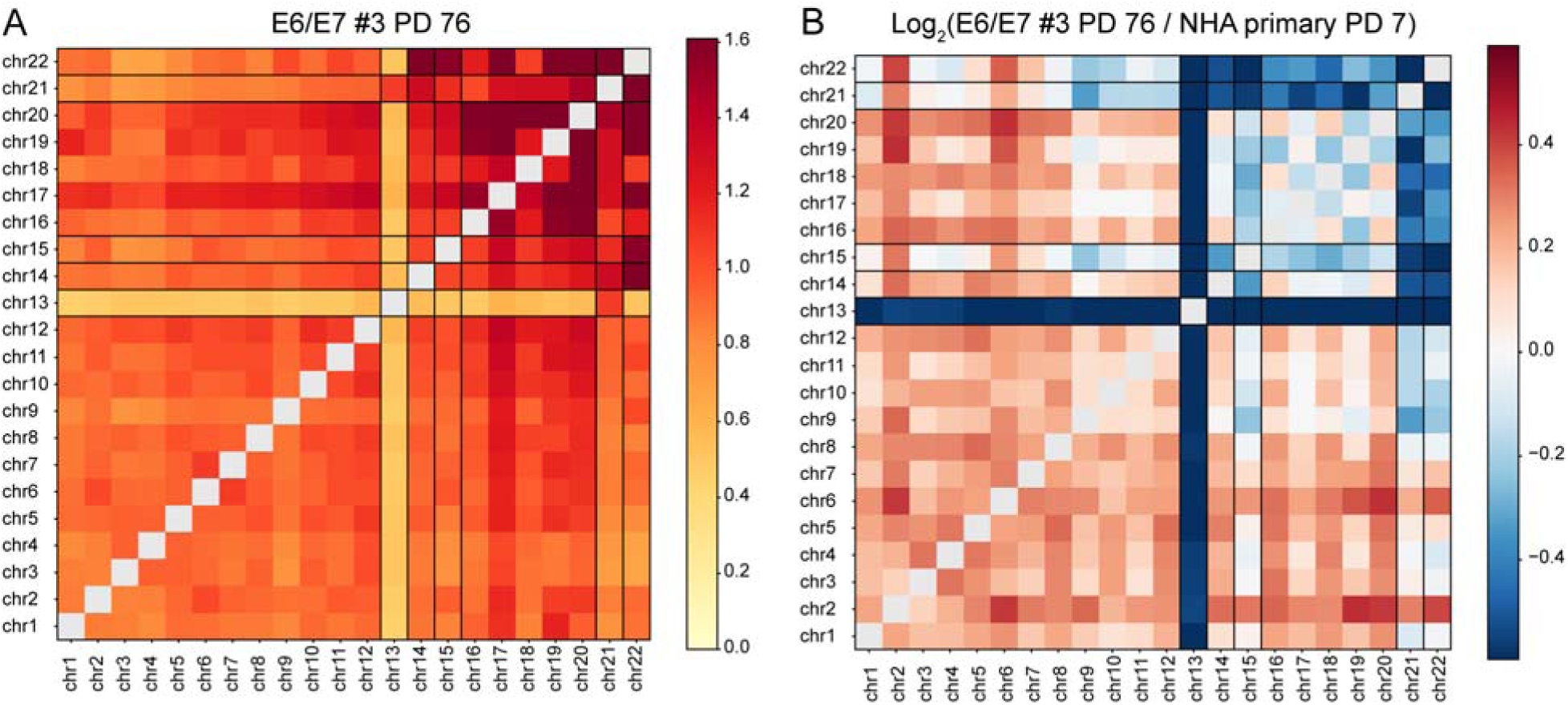
KaryoScope analysis of NHA E6/E7 #3 PD 76. (A) KaryoScope-derived inter-chromosomal contact map for NHA E6/E7 #3 PD 76, displayed as observed/expected (O/E) normalized contact frequencies. (B) Log₂FC ratio of O/E-normalized inter-chromosomal contact frequencies between NHA E6/E7 #3 PD 76 and NHA primary PD 7. Blue indicates contact depletion in E6/E7 cells, red indicates enrichment. The chromosome 13 depletion signature and altered acrocentric contacts observed at PD 48 (Figure 5E) persist at PD 76, indicating that the nucleolar reorganization phenotype is maintained across passages.

